# SARS-CoV-2 variants impact RBD conformational dynamics and ACE2 accessibility

**DOI:** 10.1101/2021.11.30.470470

**Authors:** Mariana Valério, Luís Borges-Araújo, Manuel N. Melo, Diana Lousa, Cláudio M. Soares

**Affiliations:** Instituto de Tecnologia Química e Biológica António Xavier, Universidade Nova de Lisboa, Av. da República, 2780-157 Oeiras, Portugal; iBB-Institute for Bioengineering and Biosciences, Instituto Superior Técnico, Universidade de Lisboa, Lisbon, Portugal; Associate Laboratory i4HB—Institute for Health and Bioeconomy at Instituto Superior Técnico, Universidade de Lisboa, Lisbon, Portugal

## Abstract

The coronavirus disease 2019 (COVID-19) pandemic, caused by the severe acute respiratory syndrome coronavirus 2 (SARS-CoV-2), has killed over 5 million people and is causing a devastating social and economic impact all over the world. The rise of new variants of concern (VOCs) represents a difficult challenge due to the loss vaccine and natural immunity, and increased transmissibility. All circulating VOCs contain mutations in the spike glycoprotein, which mediates fusion between the viral and host cell membranes, via its receptor binding domain (RBD) that binds to angiotensin-converting enzyme 2 (ACE2). In an attempt to understand the effect of RBD mutations in circulating VOCs, a lot of attention has been given to the RBD-ACE2 interaction. However, this type of analysis is limited, since it ignores more indirect effects, such as the conformational dynamics of the RBD itself. Observing that some VOCs mutations occur in residues that are not in direct contact with ACE2, we hypothesized that they could affect RBD conformational dynamics. To test this, we performed long atomistic (AA) molecular dynamics (MD) simulations to investigate the structural dynamics of *wt* RBD, and that of three circulating VOCs (alpha, beta, and delta). Our results show that in solution, wt RBD presents two distinct conformations: an “open” conformation where it is free to bind ACE2; and a “closed” conformation, where the RBM ridge blocks the binding surface. The alpha and beta variants significantly impact the open/closed equilibrium, shifting it towards the open conformation by roughly 20%. This shift likely increases ACE2 binding affinity. Simulations of the currently predominant delta variant RBD were extreme in this regard, in that a closed conformation was never observed. Instead, the system alternated between the before mentioned open conformation and an alternative “reversed” one, with a significantly changed orientation of the RBM ridge flanking the RBD. This alternate conformation could potentially provide a fitness advantage not only due to increased availability for ACE2 binding, but also by aiding antibody escape through epitope occlusion. These results support the hypothesis that VOCs, and particularly the delta variant, impact RBD conformational dynamics in a direction that simultaneously promotes efficient binding to ACE2 and antibody escape.

## INTRODUCTION

Coronavirus disease 2019 (COVID-19), caused by the severe acute respiratory syndrome coronavirus 2 (SARS-CoV-2)^1–3^, is a global pandemic with higher mortality than that of seasonal influenza^4^. As of November 2021, 0ver 5 million lives had been claimed by this disease^5^. Infection by SARS-CoV-2 requires the fusion of viral and host cell membranes, at either the cell surface or the endosomal membrane^6^. As for the severe acute respiratory syndrome coronavirus (SARS-CoV) and the Middle East respiratory syndrome-related coronavirus (MERS-CoV), the SARS-CoV-2 fusion process is mediated by the viral envelope spike (S) glycoprotein^6^. Upon viral attachment or uptake, host factors trigger large-scale conformational rearrangements in the S protein, including a refolding step that leads directly to membrane fusion and viral entry ^7–12^.

The SARS-CoV-2 S protein is composed of a signal peptide located at the N-terminus (residues 1-13) and 2 subunits, S1 (residues 14-685) and S2 (residues 686-1273)^13^. The S1 and S2 subunits are responsible for receptor binding and membrane fusion, respectively^13^. The S1 subunit consists of a N-terminal domain (residues 14-305) and a receptor binding domain, or RBD (residues 319-541). The RBD is responsible for the interaction of SARS-CoV-2 with host cells via binding to the angiotensin-converting enzyme 2 (ACE2)^8,10,13,14^, a regulator of the renin-angiotensin system. Binding to ACE2 is one of the first steps in what is considered to be the main mode of SARS-CoV-2 viral entry, and as such, a lot of attention has been given to the SARS-CoV-2 RBD – ACE2 complex due to both its mechanistic implications^15–20^ and pharmaceutical potential^21–27^. However, not much attention has been given to the dynamics of the RBD by itself.

The RBD core structure when bound to ACE2 (Figure 1A) consists of a twisted five stranded antiparallel β sheet (β1, β2, β3, β4 and β7), with short connecting helices and loops^28^. This core β sheet structure is further stabilized by 3 disulfide bonds. Between the core β4 and β7 strands (residues 438-506), there is an extended region containing 2 short β strands (β5 and β6), the alpha 4 and alpha 5 helices and loops. This region is the receptor-binding motif (RBM), which contains most of the residues that are responsible for interacting with ACE2^28,29^. When complexed with ACE2, the RBM folds into a concave surface, that accommodates the N-terminal α-helix of ACE2, with a ridge (residues 471 to 491) on one side, formed by a disulfide-bridge-stabilized loop (Cys480–Cys488). It is in this surface that several RBM residues establish specific and non-specific interaction with ACE2 residues^28^.

**Figure 1.**
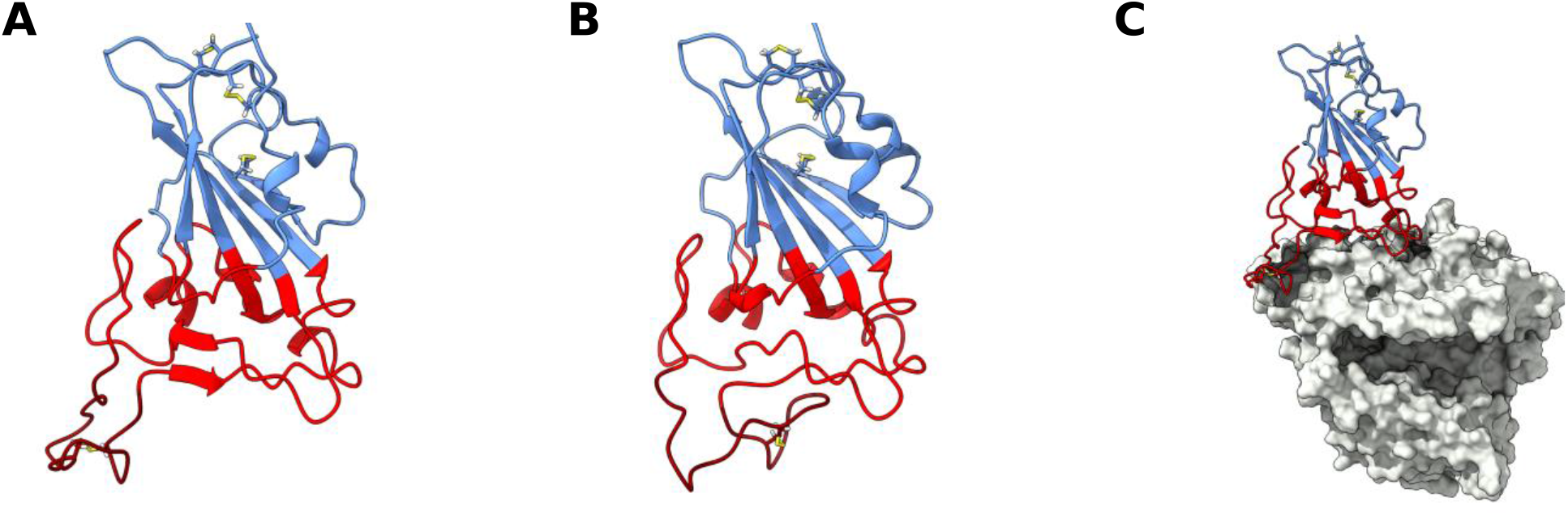
SARS-CoV-2 receptor binding domain (RBD) structure. Structure of *wt* RBD in the open (A) and closed (B) conformations. Snapshots obtained from the AA MD simulations. Disulfide bonds are represented in yellow sticks. Structure of *wt* RBD bound to ACE2 is also shown (C). The RBM region is colored red and the ridge in dark red, with the rest of the protein being colored in blue. ACE2 is in grey.

From the available experimental structural data the core β-sheet structure is very stable, but the RBM seems to be quite dynamic and not as structurally defined, unless bound to other proteins, like ACE2^14,30–33^ or antibody fragments^34–40^. Molecular dynamics (MD) simulation studies have also mostly focused on RBD complexed with these proteins, and while there are MD simulation studies of free RBD, they have either been short simulations^41–43^ or not focused on RBM dynamics^41,44^. As such, not much is known about the conformational dynamics of this motif when unbound. This is relevant because the conformational dynamics of the SARS-CoV-2 RBD and RBM might not only play an important role in receptor recognition and binding but also provide important information for the development of newer improved pharmaceuticals.

Recently, a significant number of naturally occurring mutations to the SARS-CoV-2 S protein have also been reported^45–48^. Many of these mutations have been identified in the RBD, some of which have rapidly become the dominant viral variant in certain regions due to their significant fitness advantage^45–48^. Many of these RBD mutations are thought to increase fitness by increasing ACE2 binding affinity or by escaping neutralization by anti-SARS-CoV-2 monoclonal antibodies^49^. Still, the impact of these mutations on the structural dynamics of RBD and the RBM have not yet been investigated.

In this work, we use atomistic (AA) molecular dynamics (MD) simulation methods to investigate the structural dynamics of SARS-CoV-2 RBD, and that of three naturally occurring variants of concern (VOCs): B.1.1.7, or alpha, variant^47^ (N501Y); B.1.351, or beta, variant^45^ (K417N E484K N501Y); and B.1.617.2, or delta, variant^46^ (L452R T478K). Our results show that the RBM dynamics of *wt* RBD are such that it is not always in a conformation competent for ACE2 binding (Figure 1). Conversely, all variants, and delta in particular, stabilize binding-competent configurations. The conformational space visited by the variants thus putatively increases ACE2 binding efficiency and may further provide fitness advantage by aiding in antibody escape.

## METHODS

### Molecular dynamics simulations

All atomistic simulations were performed with the GROMACS 2020.3^50,51^ package and modelled using the Amber14sb^52^, forcefield alongside the TIP3P water model^53^. The initial *wt* RBD structure was obtained from PDB ID: 6M0J^30^, which corresponds to an ACE2 bound conformation of RBD; ACE2 was excluded from this structure. The different RBD variants were generated by mutating the appropriate residues in the *wt* RBD using PyMOL^54^. Simulations were performed on each RBD protein structure in water. Each structure was inserted in a truncated dodecahedron box filled with water molecules (considering a minimum distance of 1.2 nm between protein and box walls). The total charge of the system was neutralized with the required number of Na^+^ ions, with additional Na^+^ and Cl^−^ ions added to the solution to reach an ionic strength of 0.1 M.

The system was energy-minimized using the steepest descent method for a maximum of 50000 steps with position restraints on the heteroatom positions by restraining them to the crystallographic coordinates using a force constant of 1000 kJ/mol in the X, Y and Z positions. Before performing the production runs, an initialization process was carried out in 5 stages of 100 ps each. Initially, all heavy-atoms were restrained using a force constant of 1000 kJ/mol/nm, and at the final stage only the only Cα atoms were position-restrained using the same force constant. In the first stage, the Berendsen thermostat^55^ was used to initialize and maintain the simulation at 300 K, using a temperature coupling constant of 0.01 ps, without pressure control. The second stage continued to use the Berendsen thermostat but now with a coupling constant of 0.1 ps. The third stage kept the same temperature control, but introduced isotropic pressure coupling with the Berendsen barostat^55^, with a coupling constant of 5.0 ps. The fourth stage changed the thermostat to V-rescale^56^, with a temperature coupling constant of 0.1 ps, and the barostat to Parrinello-Rahman^57^ with a pressure coupling constant of 5.0 ps. The fifth stage is equal to the fourth stage, but position restraints are only applied on Cα atoms. For production simulations, conditions were the same as for the fifth stage, but without any restraints. In all cases, 2 fs integration steps were used. Long-range electrostatic interactions were treated with the PME^58,59^ scheme, using a grid spacing of 0.12 nm, with cubic interpolation. The neighbor list was updated every twenty steps with a Verlet cutoff with a 0.8 nm radius. All bonds were constrained using the LINCS algorithm^60^.

Simulations of each of the RBD proteins were performed for at least 7 μs over 5 replicates (the *wt* was simulated for 15 μs, and the alpha, beta and delta variants for 7 μs each). The first 3 μs of simulation were considered as equilibration time and the remaining frames were used for analysis. Visualization and rendering of simulation snapshots was performed with the molecular graphics viewers VMD^61^, PyMOL^54^ and UCSF Chimera^62^.

### Principal Component Analysis

PCA is a standard dimensionality reduction method that we apply here to the (3N-6)-dimensional space of possible RBD conformations (in our case, N being the number of RBD residues). PCA consists of a linear transformation that changes a set of possibly correlated dimensions into a set of linearly uncorrelated, mutually orthogonal ones, called principal components (PCs). The first PC can be defined as the direction that accounts for as much of the variance in the data as possible, with each successive PC accounting for as much of the remaining variance as possible. Reduction of data dimensionality is achieved by retaining only a few of the first PCs — which represent the strongest correlations in the data, in our case, the most important conformational motions —, thus sacrificing some information for simplicity. Discussions of the mathematical and computational backgrounds can be found elsewhere^63–66^.

In this work, PCA was applied to sets of conformational coordinates obtained from MD simulations. Prior to PCA, each conformation was translationally and rotationally fitted to the RBD core Cα carbons of the *wt* crystal structure (hence the –6 in the dimensionality). PCs were determined using MDAnalysis^67^, from the entire pool of simulation trajectories, considering only the coordinates of the RBD’s Cα carbons. The dimensionality was reduced to the 2 most representative PCs, preserving a large part of the variance. RBD structures for each simulation frame, for each variant, could then be projected as points in this two-dimensional space, enabling a simplified visual representation of the conformation space explored by the RBD in each case.

The probability density function for each trajectory projection was estimated using a gaussian kernel estimator^65,68^ implemented in LandscapeTools’ *get_density* software as described elsewhere^65,69^. This procedure defines a probability density function P(r), with the values of P(r) being stored for the position of each data point and for the nodes of a two-dimensional uniform grid, with a mesh size of 0.5 Å. These values were used to define an energy surface, calculated as^65^:

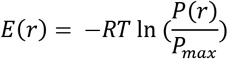

Where P_max_ is the maximum of the probability density function, P(r). The energy surface landscapes were analyzed by determining the energy minima and respective basins. The basins were defined as the set of all conformations whose steepest descent path along the energy surface leads to a particular minimum^65,70,71^. Here, the steepest descent paths for each grid cell were computed, with each conformation inheriting the path of its corresponding grid cell. Landscape regions with E > 6 k_B_T were discarded, resulting in the final set of basins for each data set.

### Residue interaction network analysis

Residue interaction networks (RINs) are graph representations of protein structures, where the nodes represent amino acid residues and the edges represent interactions between residues. Pairwise residue interactions were analyzed for the 5000 lowest energy conformations obtained for the most populated open, closed and reversed conformation basins of the energy surface landscapes of each RBD variant, using RIP-MD^72^. Several types of interactions between AAs were probed: Cα contacts, hydrogen bonds, salt bridges, disulfide bonds, cation-π, π–π, Arg-Arg, Coulomb and van der Waals. The parameters defining each interaction, as well as their mathematical formulation can be found elsewhere^72^. Once the interactions were determined, the interaction networks were visualized using Cytoscape^73^.

## RESULTS AND DISCUSSION

Our aim was to study the conformational dynamics of the SARS-CoV-2 RBD, as well as that of several other SARS-CoV-2 variants in solution. To this effect, we simulated the *wt*, alpha, beta and delta variants of the SARS-CoV-2 RBD. The gamma variant was not studied due to its similarity to the beta variant (in the RBD region, both variants share the E484K and N501Y mutations; the beta variant also contains the K417N mutation while the gamma variant has K417T).

### *wt* RBD presents two distinct RBM conformations in aqueous solution

Visual inspection of the trajectories obtained in the simulation of *wt* RBD in water revealed that large dynamic conformational changes occur in the RBM region (Figure 2A, Supplementary Video S1). The dynamics observed appear to show an opening and closing of the ACE2 binding surface of the RBM. To better characterize these conformational dynamics, we performed principal component analysis (PCA) on the coordinates recovered from these simulations, reducing them to 2 principal components; this 2D configuration space sampling was expressed as free energy landscapes (Figure 2).

**Figure 2.**
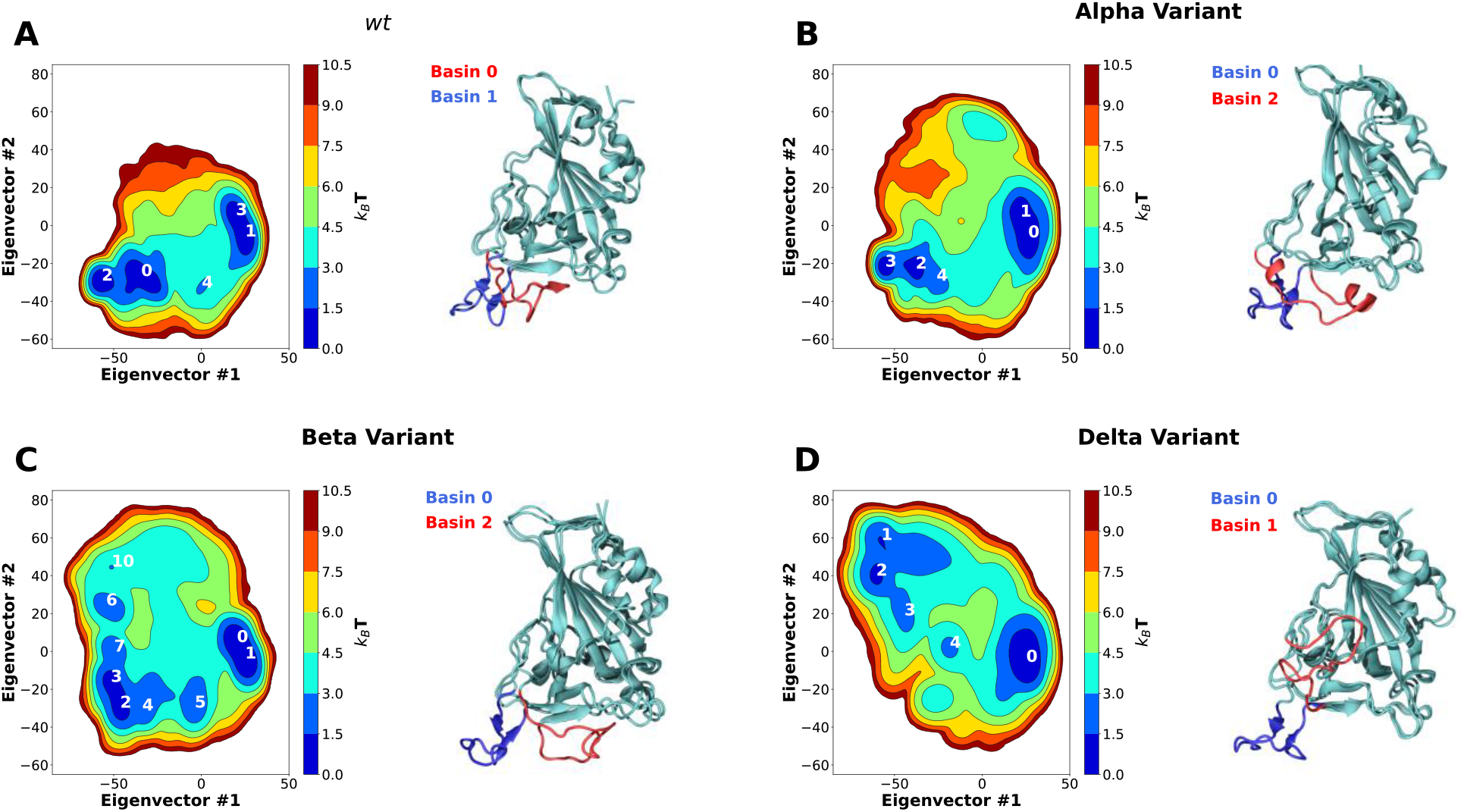
Two-dimension principal component analysis (PCA) of SARS-CoV-2 RBD conformational dynamics in water. Plots of the first two principal components determined from the Cα backbone of the *wt* RBD (A) as well as the alpha (B), beta (C) and delta (D) variants. Basins with KBT < 3 are numbered in each figure. Snapshots of the lowest energy structures for selected open and closed basins are also shown. The ridge regions of the open and closed snapshots are colored in blue and red, respectively.

For *wt* RBD, we observe two deep basin clusters (Figure 2A), as well as several other lesser basins. Closer analysis on the RBD conformations that make up each basin shows that *wt* basins 1 and 3 correspond to conformations close to the ACE2-bound one determined by X-ray crystallography^30^ (Figure 1A and 2A). We named these “open” configurations. The second basin cluster (basins 0 and 2), however, was made up by quite distinct conformations. In these basins, the loop that makes up the RBM has twisted, and collapsed over the ACE2 binding surface, effectively hiding it from the solvent (Figure 1B and 2A). We named these conformations “closed”. Further analysis of the PCA results reveals that the *wt* RBD is in a closed state for more than half of the simulation time (~55.5%, Supplementary Table S1). Given that in these conformations the RBM closes on itself, hiding the ACE2 binding surface, we can speculate that RBD would be unable to effectively bind to ACE2 and initiate the ACE2-dependent infection process. Moreover, the open and closed states were visited reversibly (Supplementary Figure S1), indicating that our simulations were not kinetically trapped in either basin.

Residue interaction network (RIN) analysis was performed for the 5000 lowest energy structures of basins 1 (open) and 0 (closed). From the identified interactions, we selected those that were present in over 50% of the simulation frames (Supplementary Figure S3). We also only considered interactions that are established by RBM residues, or those in their immediate vicinity. These RINs were then used to probe the different intramolecular interactions established in each of the conformations.

In the open conformation, the RBD ridge is stabilized by a triple π-stacking interaction between residues Y489-F456-Y473 and a hydrogen bond between Y489-Y473. Additionally, two hydrogen bonds are established between residues Y453 and E493, which help stabilize the formation of a small β-sheet (Figure 3A).

**Figure 3.**
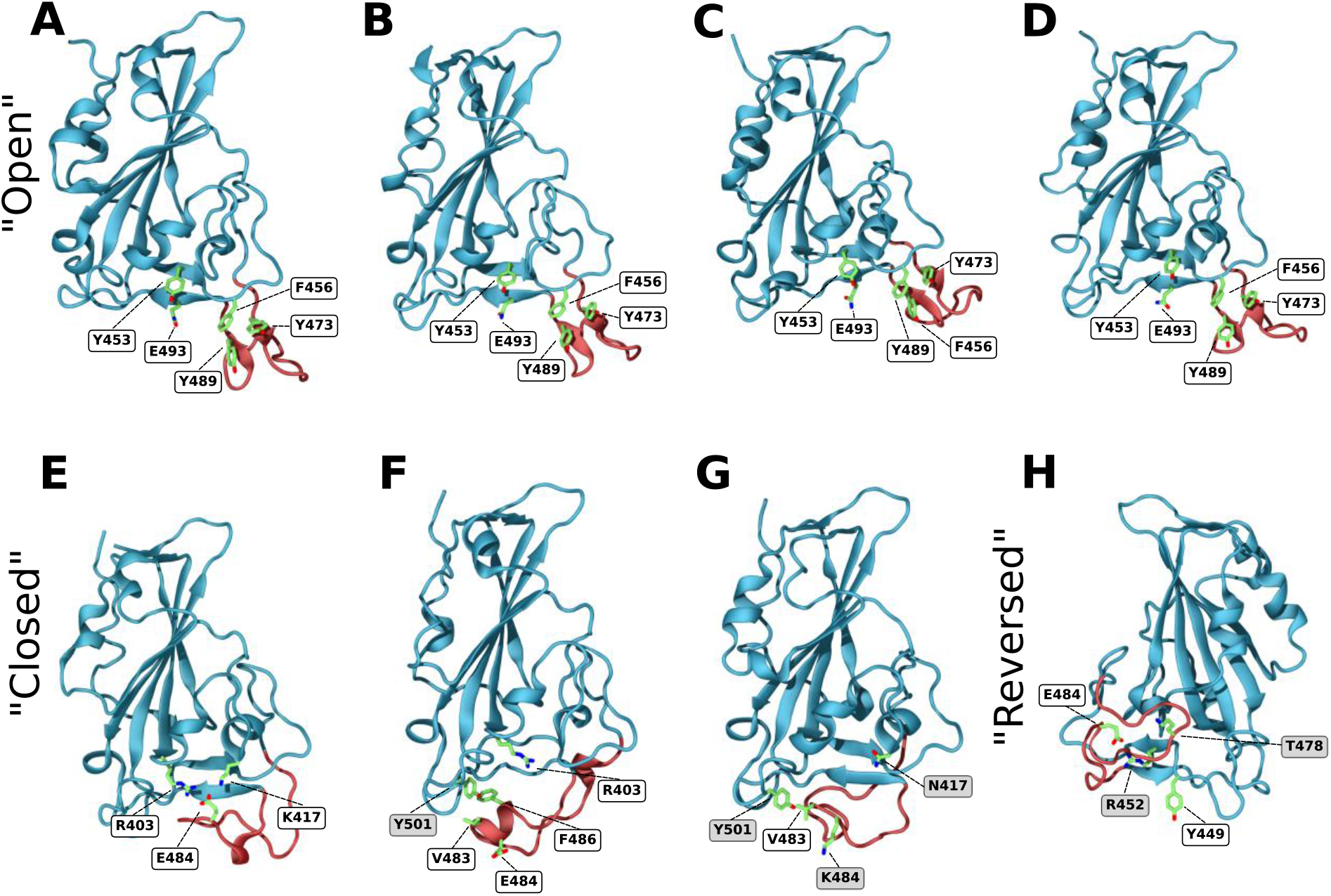
Closeup snapshots of SARS-CoV-2 RBD intramolecular interactions that stabilize the various conformations. Snapshots from AA MD simulations showcasing crucial intramolecular interactions responsible for stabilizing the open, closed, and reversed conformations for the *wt* (A, E), alpha (B, F), beta (C, G) and delta (D, H) RBD variants. The ridge region of the RBD is colored in red and residues of interest in green. Text labels indicate relevant residues, with shaded labels indicating mutations relative to *wt*. All figures are rotated 180° relative to Figures 1 and 2, apart from snapshot H.

In the closed conformation, however, the π-stacking interactions are broken, and new interactions with RBD core residues are formed in their place. F456 forms a stable π-stacking with Y421, Y489 forms a transient π-stacking interaction with F486 and Y473 forms a hydrogen bond with the backbone of Y451. Moreover, E484 forms a salt bridge with R403, that is found in the RBD core, and a hydrogen bond with K417 (Figure 3E). This hydrogen bond does not show up in the RIN, as K417 can establish a bond with each of the two glutamate oxygens, each with ~40% prevalence (each thus below our 50% selection cutoff). These two interactions, together with the formation of three hydrogen bonds (C480–S494–G482–Q493) are responsible for the “closing” of the ridge and consequent shielding of the ACE2 binding surface. The importance of the E484-R403 and E484–K417 interactions for the closing of the loop was confirmed by simulating the E484K and K417N mutants. Either of these single mutations were enough to completely deplete the closed conformation (Supplementary Figure S2A and B for E484K and K417N, respectively). This shows that both these interactions are crucial for the stabilization of the *wt* closed state. Still, several other transient hydrogen bonds, formed between residues L492, G493 and S494 of strand β6, and T478, C480, N481, G482 and E484 of the RBM ridge, assist in stabilizing the structure.

The closed conformation does not seem to substantially impact RBD secondary structure (Supplementary Figure S7). The largest impact appears to be limited to residues 473-474 and 488-489, that in the open state display a slight β-sheet character. However, upon closing, this β-sheet character disappears. This effect comes from residues 473 and 489 no longer participating in the triple π-stacking that was likely stabilizing this region.

Apart from impacting ACE2 accessibility, the closing of the RBM ridge also decreases the solvent accessible surface area (SASA) of the RBD by slightly over 3 % (Supplementary Table S2).

Although other studies have noted the high flexibility in the RBM region of the RBD^41–43^, this is, as far as we know, the first report of this “hinge” mechanism which can effectively hide the ACE2 binding surface of the RBD from binding partners. While it is likely that induced fit interactions might assist in opening a closed conformation for binding to ACE2, it is safe to assume that the closed conformation will have its binding to ACE2 substantially hindered when compared to an open conformation.

### SARS-CoV-2 alpha and beta variants impact RBM conformational dynamics and exposure

The first SARS-CoV-2 variant of concern to be identified was first detected in the UK. It is often referred to as B.1.1.7 or alpha variant and has only one mutation in the RBD region — N501Y. A second variant emerged soon after in South Africa, independently of B.1.1.7, referred to as B.1.351 or beta variant. In the RBD region, this variant shares the N501Y mutation with the alpha variant and includes two others: K417N and E484K^47^.

Like the *wt* variant, MD simulations of the RBDs from the alpha and beta variants also showed the prevalence of two sets of RBM conformations, corresponding to open and closed conformations (Supplementary Videos S2 and S3). PCA analysis of the alpha variant trajectory shows two deep basin clusters (Figure 2B), basins 0 and 1, and basins 2 and 3, which correspond to open and closed conformations respectively. However, unlike the *wt* variant, the alpha variant remains most of the simulation time in an open conformation (~72.64 %, Supplementary Table S1). The beta variant (Figure 2C) also has two deep basin clusters (basins 0 and 1, and basins 2 and 3), corresponding to open and closed conformations, respectively. Like the alpha variant, beta remains in an open conformation for substantially longer time than the *wt* (~69 %, Supplementary Table S1). In both cases, and as for *wt*, our simulations were able to reversibly visit either state (Supplementary Figure S1).

Both alpha and beta variants shift the open/closed equilibrium towards more open conformations by roughly 20%. A closing ΔΔG was calculated from the ratio between time spent in the open and closed states, where the time spent in each individual open and closed basin was added together (Supplementary Table S1). The equilibrium shift led to a decrease in the closing ΔΔG from 0.55 ± 0.17 kJ/mol, in the case of *wt* RBD, to −2.44 ± 0.22 and −2.09 ± 0.14 kJ/mol, for the alpha and beta variants, respectively. As mentioned previously, it is likely that only the open conformations are fully available to bind to ACE2, meaning that these mutations substantially increase the accessibility of RBD to ACE2, and probably impact ACE2-RBD binding.

By analyzing the intramolecular residue interactions for both variants we observe that the interactions which stabilize the open conformation in the *wt* variant are conserved in both alpha and beta variants, namely the triple π-stacking between residues Y489–F456–Y473, as well as the hydrogen bond between Y489 and Y473. An additional hydrogen bond between Q493 and Y453 assists in stabilizing the β6 strand (Figure 3B and 3C).

Interestingly, in both the open and closed conformations of the alpha variant, the interactions established by residue Y501 (alpha’s only mutation in the RBD) that were previously present in the *wt* variant are maintained in the alpha variant (two hydrogen bonds established through the residue backbones: Q458–Y501 and Y501–Q506). However, the main interactions that stabilize the closed conformations differ between the alpha variant and *wt* (although some transient hydrogen bonds between strand β6 and the RBM ridge do remain). Instead of the E484–R403 salt bridge seen for *wt*, in the alpha variant the closed conformation is promoted by the formation of hydrophobic interactions between the mutated Y501, V483 and F486 (Figure 3F). This arrangement hinders the establishment of the E484–R403 salt-bridge (as can be seen in Supplementary Video 2) while being itself less stable than the open conformations. This is the likely cause for the decrease in percentage of closed state observed for alpha. Progression to the E484–R403 salt-bridge may also be prevented in part by the establishment of a short α-helix, discussed ahead.

In the beta variant, the closed conformation is notably impacted by both the E484K and the N501Y mutations. The E484K mutation prevents the formation of the E484–R403 salt bridge that was crucial for the stability of the closed conformation in the *wt* variant. However, unlike the single E484K mutant (Supplementary Figure S2), the beta variant can still reach a closed conformation. This is because it can establish the same hydrophobic interaction between Y501 and V483 as the alpha variant (Figure 3G). This closed state is also stabilized by the same transient hydrogen bonds between strand β6 and the RBM ridge seen in the *wt* and alpha variants Concerning the secondary structure, there are no substantial differences between the alpha or beta open states and the *wt* open state (Figure S7). However, upon closing, both alpha and delta form a small α-helix between residues 475 and 490, for roughly 30% of the simulation time. This helical character might be relevant for the alpha variant, as it assists in facing the E484 sidechain away from R403 (Figure 3F and 3G), hindering the formation of the salt-bridge. Additionally, the alpha variant also shows some helicity in residues 482 to 489, which likely arises from contacts between residues in this helix and the mutated N501Y.

Curiously, while the alpha variant also shows a considerable decrease in SASA upon closing (~5%), the beta variant shows no substantial change.

Overall, these results showcase a possible alternative mechanism for how the alpha and beta variants might facilitate viral entry into the host cells. By shifting the open/closed equilibrium towards the ACE2-accessible open conformation, both of these variants are facilitating ACE2–RBD binding, which will inevitably lead to an increase in binding affinity and enhanced receptor-dependent infection.

### SARS-CoV-2 delta variant shows conformational dynamics distinct from the other variants

As of November 2021 the global dominant SARS-CoV-2 variant is the B.1.617.2 (or delta)^46^. It contains two mutations in the RBD region: L452R and T478K. Like the *wt*, alpha and beta variants, MD simulations of the delta RBD show the prevalence of two sets of RBM conformations, one of which corresponds to the *wt* open conformation (Supplementary Video S4) and is stabilized by the same interactions observed for the three other variants (Figure 3D). However, unlike those variants, MD simulations of the delta RBD do not show the occurrence of a closed conformation at all. Instead, an alternative open conformation is present, which we refer to as reversed. PCA analysis of the delta variant trajectory, show two deep basins, 0 and 2 in Figure 2D, which correspond to the open and reversed conformations, respectively. As for the other variants, simulations were able to reversibly visit the two states (Supplementary Figure S1).

The reversed conformation showcases the incredible flexibility of the RBM region, which not only opens and closes over the ACE2 binding surface of the RBD but acts as a two-way hinge that reverse-folds to the side of the RBD. This alternative conformation might also prove significant advantages over the *wt* open state: RBD-targeting antibodies are known to bind via recognition of the RBM ridge region^17,74^; the reversed state putatively hides this region from antibody recognition, while still providing an open ACE2 binding surface for infection.

A hydrogen bond between the mutated R452 on strand β5 and Y449 appears to be one of the main driving forces folding the delta variant’s ridge region backwards. This interaction destabilizes the β5 strand and enables the ridge to move up and interact with the core. Transient interactions between ridge residues G476, S477 as well as the mutated K478 with residues R346, F347 and N354 of strand β1 stabilize the contact between the ridge loop and the RBD core, keeping it locked in place (Figure 3H).

Regarding the secondary structure, much like the other variants, the delta open conformation is very similar to that of the *wt* (Supplementary Table S3). However, as expected, the reversed conformation shows substantial differences. In this state, the two small beta strands formed by residues 473-474 and 488-489, present in the open conformation, are completely lost. Additionally, the beta-sheet formed by strands β5 and β6 becomes less prevalent, likely due to the L452R mutation (one of the β5 strand residues that destabilizes the β-sheet by establishing a new interaction with Y449). Curiously, like in the alpha and beta variants, there is also a significant alpha helical character between residues 490 and 475.

As for the closed conformations of the *wt*, alpha and beta variants, the delta reversed conformation also leads to a decrease in SASA (~ 3%). Unlike the closed conformations, however, this alternative open conformation still presents a fully accessible ACE2 binding surface.

### Impact of SARS-CoV-2 variants on ACE2 binding affinity

To find experimental basis for our results, we compiled ACE2-RBD binding kinetics data from recent studies^75–81^ (Supplementary Table S3). These results were obtained by surface plasmon resonance (SPR) and biolayer interferometry (BLI) and encompass data regarding both the *wt* and studied variants. Additionally, we compiled results obtained for just the RBD as well as for the entire S protein. While the binding kinetics values recovered from these studies are not fully consistent with each other, likely due to differences in particular experimental setups, they are mostly in the same range, and appear to follow similar trends.

Regarding the equilibrium dissociation constant (*K*_*d*_), all variants have an increased binding affinity when compared to the *wt*. With the currently available data, however, it is hard to distinguish between the efficiency of the several variants, with the alpha and beta variants showing a slightly better affinity than delta.

To get more information, we analyzed both the association (*k*_*on*_) and dissociation rate constants (*k*_*off*_). *k*_*off*_ reflects the lifetime of the protein-protein complex and as such, the strength of the interaction. We observe a consistent decrease in *k*_*off*_ for the variants in comparison to the *wt*. The alpha and beta variants stand out from delta in this regard, with substantially lower *k*_*off*_ values. These results hint at the variants interacting more strongly with ACE2 than the *wt*, with the alpha and beta complexes being substantially more stable than those of delta. Several other MD studies have studied the impact of these mutations on the contacts between RBD and ACE2 and have shown how the substantially altered ACE2-RBD interaction network of the alpha and beta variants might be outperforming that of the *wt* variant^82–86^. The delta variant does not contain mutations to the RBD ACE2 binding surface and, as such, the interactions established are not substantially different from those of *wt*. This is reflected in a closer, although still lower, *k*_*off*_ value.

The variants also substantially impact *k*_*on*_. This rate constant reflects the efficiency with which protein–protein collisions lead to a bound state. While a couple of studies show no significant impact^75,79^, most show that the variants lead to a substantial increase in *k*_*on*_, reflecting an increase in ACE2 accessibility^76–78,80,81^. We propose that this can be explained by the significant changes in RBM conformational dynamics that we have here described, where mutations lead to a decrease in prevalence of the closed state, favoring binding. As such, our results point to an alternative mechanism for enhancing RBD-ACE2 binding, not by directly strengthening ACE2-RBD interactions, but rather by boosting, via modulation of ridge dynamics, the ACE2 binding competence.

### Emergent VOCs share relevant RBM mutations with alpha beta and delta

Recently, a new VOC — B.1.1.529 or omicron — has emerged which is overtaking delta as the dominant variant in some world regions^87^. The omicron variant contains 15 mutations in the RBD region, 10 of which are concentrated in the RBM. Some of those mutations are also observed, or are similar to those, in the alpha, beta and delta variants: K417N, T478K, E484A and N501Y. From our work, we can expect this large number of mutations to heavily impact the open/closed equilibrium we observed for *wt* RBD. In particular, the presence of the T478K mutation — shared with the delta variant — points towards possible alternative conformations like delta’s reversed state. Just as for delta, these conformations are likely to improve antibody escape, providing omicron with a substantial fitness advantage.

## CONCLUSION

In this work we performed AA MD simulations of the SARS-CoV-2 RBD, as well as that of the alpha, beta and delta VOCs, to characterize the impact of the mutations on RBD conformational dynamics in solution.

Our results show that the *wt* RBD adopts two distinct conformations in equilibrium: an open conformation where the RBD is free to bind ACE2; and a closed conformation, where the RBM ridge blocks the ACE2 binding surface and likely hinders binding to ACE2. We characterized the two states and showed that they originate from specific intramolecular interactions between residues of the RBM ridge and those of the ACE2 binding surface. As far as we know, this is the first report of this “hinge-like” mechanism which can effectively shield the ACE2 binding surface from the solvent and binding partners. This mechanism is yet to be seen in experimentally solved RBD structures, which have thus far struggled to fully resolve the unbound RBM region^20,29,88^. The RBM is found unresolved in most structures due to the large flexibility of the region, and those that are fully resolved are often structures of RBD complexed with either ACE2^14,30–33^, antibodies^34–40^ or itself by dimerizing via the ACE2 binding surface^89,90^.

The three variants tested in this work, significantly impacted the open/closed equilibrium we observed for *wt* RBD. Both alpha and beta variants shifted the equilibrium towards more open conformations by roughly 20%, while the delta variant did not show the presence of a closed conformation at all. This shift towards more open conformations likely enhances ACE2 binding affinity by increasing accessibility to the RBM and facilitating binding. Several experimental binding studies have shown that these variants lead to a substantial increase in ACE2-RBD binding association rate constant, reflecting an increased ACE2 accessibility, in agreement with our findings.

Additionally, the delta variant showed an alternative open conformation, distinct from that of the other variants.

This alternative conformation keeps the ACE2 binding surface open and accessible for binding, but significantly alters the conformation of the RBM ridge. This state presents a substantially altered ridge region, which bends backwards towards the RBD core, shielding some of it from exposure. We hypothesize that this may provide a fitness advantage by aiding in antibody escape: many RBD-targeting antibodies are known to target the RBM ridge region^35,74,91,92^. In the alternative open conformation, the ridge may be not as easily recognized, while the ACE2 binding surface remains unobstructed for infection.

These results show that the mutations found in the three VOCs (alpha, beta and delta) impact RBD conformational dynamics in a direction that promotes efficient binding to ACE2 and (in the case of the delta variant) antibody escape, an effect which has thus far been disregarded. In this context, our findings can also help explain some of the antibody-evading characteristics of the emergent omicron variant.

## ASSOCIATED CONTENT

### Supporting Information

Extended materials providing video description, additional analysis results on RMSD, SASA, RINs, additional detail on the PCA analysis and compilation of ACE2-RBD experimental binding results. Videos S1 through S4 show the RBD conformational dynamics of the *wt*, alpha, beta and delta variants.

## AUTHOR INFORMATION

### Author Contributions

§ M.V. and L.B.A. contributed equally to this work. D.L. and C.M.S. designed the simulation setup. DL prepared the systems’ topologies and C.M.S. performed the simulations. M.V., L.B.A., M.N.M., D.L. and C.M.S. designed the analysis and M.V. and L.B.A. performed them. All authors contributed to manuscript writing and revision and have given approval to the final version of the manuscript.

### Notes

The authors declare no competing financial interest.

## ACKNOWLEDGMENT

The authors thank António M. Baptista and Sara R. R. Campos for their helpful discussion and input, and for making available their density and energy landscape analysis package, LandscapeTools. L.B.A. thanks the Medical Biochemistry and Biophysics Doctoral Program (M2B-PhD) and Fundação para a Ciência e a Tecnologia, I.P. (FCT) for PhD fellowship PD/BD/137492/2018. M.V. thanks FCT for the PhD fellowship SFRH/BD/148542/2019. M.N.M thanks FCT for fellowship CEECIND/04124/2017. D.L. acknowledges FCT project PTDC/CCI-BIO/28200/2017. C.M.S. and M.N.M further acknowledge FCT project MOSTMICRO-ITQB, with references UIDB/04612/2020 and UIDP/04612/2020.

## Insert Table of Contents artwork here

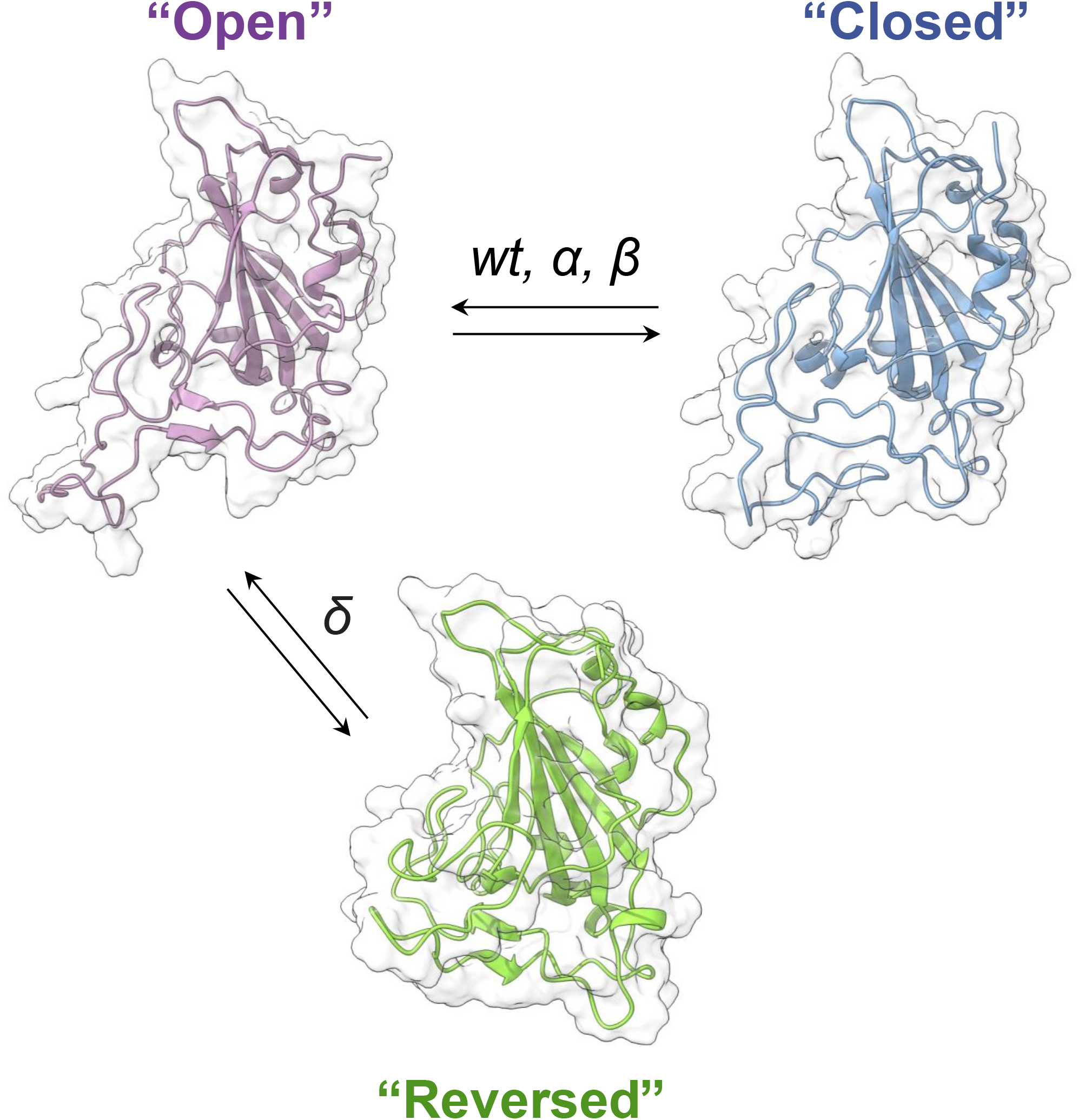

## REFERENCES

(1) Andersen, K. G.; Rambaut, A.; Lipkin, W. I.; Holmes, E. C.; Garry, R. F. The Proximal Origin of SARS-CoV-2. Nature Medicine. Nature Research April 1, 2020, pp 450–452.

(2) Wu, F.; Zhao, S.; Yu, B.; Chen, Y. M.; Wang, W.; Song, Z. G.; Hu, Y.; Tao, Z. W.; Tian, J. H.; Pei, Y. Y.; Yuan, M. L.; Zhang, Y. L.; Dai, F. H.; Liu, Y.; Wang, Q. M.; Zheng, J. J.; Xu, L.; Holmes, E. C.; Zhang, Y. Z. A New Coronavirus Associated with Human Respiratory Disease in China. Nature 2020, 579 (7798), 265–269.

(3) Zhu, N.; Zhang, D.; Wang, W.; Li, X.; Yang, B.; Song, J.; Zhao, X.; Huang, B.; Shi, W.; Lu, R.; Niu, P.; Zhan, F.; Ma, X.; Wang, D.; Xu, W.; Wu, G.; Gao, G. F.; Tan, W. A Novel Coronavirus from Patients with Pneumonia in China, 2019. N. Engl. J. Med. 2020, 382 (8), 727–733.

(4) Zhou, F.; Yu, T.; Du, R.; Fan, G.; Liu, Y.; Liu, Z.; Xiang, J.; Wang, Y.; Song, B.; Gu, X.; Guan, L.; Wei, Y.; Li, H.; Wu, X.; Xu, J.; Tu, S.; Zhang, Y.; Chen, H.; Cao, B. Clinical Course and Risk Factors for Mortality of Adult Inpatients with COVID-19 in Wuhan, China: A Retrospective Cohort Study. Lancet 2020, 395 (10229), 1054–1062.

(5) World Health Organization. WHO Coronavirus (COVID-19) Dashboard https://covid19.who.int/ (accessed Nov 29, 2021).

(6) Jackson, C. B.; Farzan, M.; Chen, B.; Choe, H. Mechanisms of SARS-CoV-2 Entry into Cells. Nat. Rev. Mol. Cell Biol. 2021, 1–18.

(7) Li, F. Structure, Function, and Evolution of Coronavirus Spike Proteins. Annual Review of Virology. Annual Reviews Inc. September 29, 2016, pp 237–261.

(8) Hoffmann, M.; Kleine-Weber, H.; Schroeder, S.; Krüger, N.; Herrler, T.; Erichsen, S.; Schiergens, T. S.; Herrler, G.; Wu, N. H.; Nitsche, A.; Müller, M. A.; Drosten, C.; Pöhlmann, S. SARSCoV-2 Cell Entry Depends on ACE2 and TMPRSS2 and Is Blocked by a Clinically Proven Protease Inhibitor. Cell 2020, 181 (2), 271–280.e8.

(9) Wan, Y.; Shang, J.; Graham, R.; Baric, R. S.; Li, F. Receptor Recognition by the Novel Coronavirus from Wuhan: An Analysis Based on Decade-Long Structural Studies of SARS Coronavirus. J. Virol. 2020, 94 (7).

(10) Bosch, B. J.; van der Zee, R.; de Haan, C. A. M.; Rottier, P. J. M. The Coronavirus Spike Protein Is a Class I Virus Fusion Protein: Structural and Functional Characterization of the Fusion Core Complex. J. Virol. 2003, 77 (16), 8801–8811.

(11) Walls, A. C.; Tortorici, M. A.; Snijder, J.; Xiong, X.; Bosch, B. J.; Rey, F. A.; Veesler, D. Tectonic Conformational Changes of a Coronavirus Spike Glycoprotein Promote Membrane Fusion. Proc. Natl. Acad. Sci. U. S. A. 2017, 114 (42), 11157–11162.

(12) de Vries, R. D.; Schmitz, K. S.; Bovier, F. T.; Predella, C.; Khao, J.; Noack, D.; Haagmans, B. L.; Herfst, S.; Stearns, K. N.; Drew-Bear, J.; Biswas, S.; Rockx, B.; McGill, G.; Dorrello, N. V.; Gellman, S. H.; Alabi, C. A.; de Swart, R. L.; Moscona, A.; Porotto, M. Intranasal Fusion Inhibitory Lipopeptide Prevents Direct-Contact SARS-CoV-2 Transmission in Ferrets. Science (80-. ). 2021, 371 (6536), eabf4896.

(13) Huang, Y.; Yang, C.; Xu, X. feng; Xu, W.; Liu, S. wen. Structural and Functional Properties of SARS-CoV-2 Spike Protein: Potential Antivirus Drug Development for COVID-19. Acta Pharmacologica Sinica. Springer Nature September 1, 2020, pp 1141–1149.

(14) Wang, Q.; Zhang, Y.; Wu, L.; Niu, S.; Song, C.; Zhang, Z.; Lu, G.; Qiao, C.; Hu, Y.; Yuen, K. Y.; Wang, Q.; Zhou, H.; Yan, J.; Qi, J. Structural and Functional Basis of SARS-CoV-2 Entry by Using Human ACE2. Cell 2020, 181 (4), 894–904.e9.

(15) Hussain, M.; Jabeen, N.; Raza, F.; Shabbir, S.; Baig, A. A.; Amanullah, A.; Aziz, B. Structural Variations in Human ACE2 May Influence Its Binding with SARS-CoV-2 Spike Protein. J. Med. Virol. 2020, 92 (9), 1580–1586.

(16) Ali, F.; Elserafy, M.; Alkordi, M. H.; Amin, M. ACE2 Coding Variants in Different Populations and Their Potential Impact on SARS-CoV-2 Binding Affinity. Biochem. Biophys. Reports 2020, 24, 100798.

(17) Alenquer, M.; Ferreira, F.; Lousa, D.; Valério, M.; Medina-Lopes, M.; Bergman, M.-L.; Gonçalves, J.; Demengeot, J.; Leite, R. B.; Lilue, J.; Ning, Z.; Penha-Gonçalves, C.; Soares, H.; Soares, C. M.; Amorim, M. J. Signatures in SARS-CoV-2 Spike Protein Conferring Escape to Neutralizing Antibodies. PLOS Pathog. 2021, 17 (8), e1009772.

(18) Lupala, C. S.; Li, X.; Lei, J.; Chen, H.; Qi, J.; Liu, H.; Su, X.-D. Computational Simulations Reveal the Binding Dynamics between Human ACE2 and the Receptor Binding Domain of SARS-CoV-2 Spike Protein. Quant. Biol. 2021, 9 (1), 61–72.

(19) Yan, F.-F.; Gao, F. Comparison of the Binding Characteristics of SARS-CoV and SARS-CoV-2 RBDs to ACE2 at Different Temperatures by MD Simulations. Brief. Bioinform. 2021, 22 (2), 1122–1136.

(20) Xu, C.; Wang, Y.; Liu, C.; Zhang, C.; Han, W.; Hong, X.; Wang, Y.; Hong, Q.; Wang, S.; Zhao, Q.; Wang, Y.; Yang, Y.; Chen, K.; Zheng, W.; Kong, L.; Wang, F.; Zuo, Q.; Huang, Z.; Cong, Y. Conformational Dynamics of SARS-CoV-2 Trimeric Spike Glycoprotein in Complex with Receptor ACE2 Revealed by Cryo-EM. Sci. Adv. 2021, 7 (1).

(21) Cao, L.; Goreshnik, I.; Coventry, B.; Case, J. B.; Miller, L.; Kozodoy, L.; Chen, R. E.; Carter, L.; Walls, A. C.; Park, Y. J.; Strauch, E. M.; Stewart, L.; Diamond, M. S.; Veesler, D.; Baker, D. De Novo Design of Picomolar SARS-CoV-2 Miniprotein Inhibitors. Science (80-. ). 2020, 370 (6515).

(22) Alexpandi, R.; De Mesquita, J. F.; Pandian, S. K.; Ravi, A. V. Quinolines-Based SARS-CoV-2 3CLpro and RdRp Inhibitors and Spike-RBD-ACE2 Inhibitor for Drug-Repurposing Against COVID-19: An in Silico Analysis. Front. Microbiol. 2020, 0, 1796.

(23) Awad, I. E.; Abu-Saleh, A. A.-A. A.; Sharma, S.; Yadav, A.; Poirier, R. A. High-Throughput Virtual Screening of Drug Databanks for Potential Inhibitors of SARS-CoV-2 Spike Glycoprotein. https://doi.org/10.1080/07391102.2020.1835721 2020.

(24) Padhi, A. K.; Seal, A.; Khan, J. M.; Ahamed, M.; Tripathi, T. Unraveling the Mechanism of Arbidol Binding and Inhibition of SARS-CoV-2: Insights from Atomistic Simulations. Eur. J. Pharmacol. 2021, 894, 173836.

(25) Kumar, V.; Liu, H.; Wu, C. Drug Repurposing against SARSCoV-2 Receptor Binding Domain Using Ensemble-Based Virtual Screening and Molecular Dynamics Simulations. Comput. Biol. Med. 2021, 135, 104634.

(26) Patel, C. N.; Goswami, D.; Jaiswal, D. G.; Parmar, R. M.; Solanki, H. A.; Pandya, H. A. Pinpointing the Potential Hits for Hindering Interaction of SARS-CoV-2 S-Protein with ACE2 from the Pool of Antiviral Phytochemicals Utilizing Molecular Docking and Molecular Dynamics (MD) Simulations. J. Mol. Graph. Model. 2021, 105, 107874.

(27) Muhseen, Z. T.; Hameed, A. R.; Al-Hasani, H. M. H.; Tahir ul Qamar, M.; Li, G. Promising Terpenes as SARS-CoV-2 Spike Receptor-Binding Domain (RBD) Attachment Inhibitors to the Human ACE2 Receptor: Integrated Computational Approach. J. Mol. Liq. 2020, 320, 114493.

(28) Lan, J.; Ge, J.; Yu, J.; Shan, S.; Zhou, H.; Fan, S.; Zhang, Q.; Shi, X.; Wang, Q.; Zhang, L.; Wang, X. Structure of the SARS-CoV-2 Spike Receptor-Binding Domain Bound to the ACE2 Receptor. Nature 2020, 581 (7807), 215–220.

(29) Wrapp, D.; Wang, N.; Corbett, K. S.; Goldsmith, J. A.; Hsieh, C.-L.; Abiona, O.; Graham, B. S.; McLellan, J. S. Cryo-EM Structure of the 2019-NCoV Spike in the Prefusion Conformation. Science (80-. ). 2020, 367 (6483), 1260–1263.

(30) Lan, J.; Ge, J.; Yu, J.; Shan, S.; Zhou, H.; Fan, S.; Zhang, Q.; Shi, X.; Wang, Q.; Zhang, L.; Wang, X. Structure of the SARS-CoV-2 Spike Receptor-Binding Domain Bound to the ACE2 Receptor. Nat. 2020 5817807 2020, 581 (7807), 215–220.

(31) Li, F. Structure of SARS Coronavirus Spike Receptor-Binding Domain Complexed with Receptor. Science (80-. ). 2005, 309 (5742), 1864–1868.

(32) Shang, J.; Ye, G.; Shi, K.; Wan, Y.; Luo, C.; Aihara, H.; Geng, Q.; Auerbach, A.; Li, F. Structural Basis of Receptor Recognition by SARS-CoV-2. Nat. 2020 5817807 2020, 581 (7807), 221–224.

(33) Zhou, T.; Tsybovsky, Y.; Gorman, J.; Rapp, M.; Cerutti, G.; Chuang, G.-Y.; Katsamba, P. S.; Sampson, J. M.; Schön, A.; Bimela, J.; Boyington, J. C.; Nazzari, A.; Olia, A. S.; Shi, W.; Sastry, M.; Stephens, T.; Stuckey, J.; Teng, I.-T.; Wang, P.; Wang, S.; Zhang, B.; Friesner, R. A.; Ho, D. D.; Mascola, J. R.; Shapiro, L.; Kwong, P. D. Cryo-EM Structures of SARS-CoV-2 Spike without and with ACE2 Reveal a PH-Dependent Switch to Mediate Endosomal Positioning of Receptor-Binding Domains. Cell Host Microbe 2020, 28 (6), 867–879.e5.

(34) Rapp, M.; Guo, Y.; Reddem, E. R.; Yu, J.; Liu, L.; Wang, P.; Cerutti, G.; Katsamba, P.; Bimela, J. S.; Bahna, F. A.; Mannepalli, S. M.; Zhang, B.; Kwong, P. D.; Huang, Y.; Ho, D. D.; Shapiro, L.; Sheng, Z. Modular Basis for Potent SARS-CoV-2 Neutralization by a Prevalent VH1-2-Derived Antibody Class. Cell Rep. 2021, 35 (1).

(35) Tortorici, M. A.; Beltramello, M.; Lempp, F. A.; Pinto, D.; Dang, H. V.; Rosen, L. E.; McCallum, M.; Bowen, J.; Minola, A.; Jaconi, S.; Zatta, F.; De Marco, A.; Guarino, B.; Bianchi, S.; Lauron, E. J.; Tucker, H.; Zhou, J.; Peter, A.; Havenar-Daughton, C.; Wojcechowskyj, J. A.; Case, J. B.; Chen, R. E.; Kaiser, H.; Montiel-Ruiz, M.; Meury, M.; Czudnochowski, N.; Spreafico, R.; Dillen, J.; Ng, C.; Sprugasci, N.; Culap, K.; Benigni, F.; Abdelnabi, R.; Foo, S. Y. C.; Schmid, M. A.; Cameroni, E.; Riva, A.; Gabrieli, A.; Galli, M.; Pizzuto, M. S.; Neyts, J.; Diamond, M. S.; Virgin, H. W.; Snell, G.; Corti, D.; Fink, K.; Veesler, D. Ultrapotent Human Antibodies Protect against SARS-CoV-2 Challenge via Multiple Mechanisms. Science (80-. ). 2020, 370 (6519), 950–957.

(36) Wu, Y.; Wang, F.; Shen, C.; Peng, W.; Li, D.; Zhao, C.; Li, Z.; Li, S.; Bi, Y.; Yang, Y.; Gong, Y.; Xiao, H.; Fan, Z.; Tan, S.; Wu, G.; Tan, W.; Lu, X.; Fan, C.; Wang, Q.; Liu, Y.; Zhang, C.; Qi, J.; Gao, G. F.; Gao, F.; Liu, L. A Noncompeting Pair of Human Neutralizing Antibodies Block COVID-19 Virus Binding to Its Receptor ACE2. Science (80-. ). 2020, 368 (6496), 1274–1278.

(37) Bertoglio, F.; Fühner, V.; Ruschig, M.; Heine, P. A.; Abassi, L.; Klünemann, T.; Rand, U.; Meier, D.; Langreder, N.; Steinke, S.; Ballmann, R.; Schneider, K.-T.; Roth, K. D. R.; Kuhn, P.; Riese, P.; Schäckermann, D.; Korn, J.; Koch, A.; Chaudhry, M. Z.; Eschke, K.; Kim, Y.; Zock-Emmenthal, S.; Becker, M.; Scholz, M.; Moreira, G. M. S. G.; Wenzel, E. V.; Russo, G.; Garritsen, H. S. P.; Casu, S.; Gerstner, A.; Roth, G.; Adler, J.; Trimpert, J.; Hermann, A.; Schirrmann, T.; Dübel, S.; Frenzel, A.; Heuvel, J. Van den; Čičin-Šain, L.; Schubert, M.; Hust, M. A SARS-CoV-2 Neutralizing Antibody Selected from COVID-19 Patients Binds to the ACE2-RBD Interface and Is Tolerant to Most Known RBD Mutations. Cell Rep. 2021, 36 (4).

(38) Kreye, J.; Reincke, S. M.; Kornau, H.-C.; Sánchez-Sendin, E.; Corman, V. M.; Liu, H.; Yuan, M.; Wu, N. C.; Zhu, X.; Lee, C.-C. D.; Trimpert, J.; Höltje, M.; Dietert, K.; Stöffler, L.; Wardenburg, N. von; Hoof, S. van; Homeyer, M. A.; Hoffmann, J.; Abdelgawad, A.; Gruber, A. D.; Bertzbach, L. D.; Vladimirova, D.; Li, L. Y.; Barthel, P. C.; Skriner, K.; Hocke, A. C.; Hippenstiel, S.; Witzenrath, M.; Suttorp, N.; Kurth, F.; Franke, C.; Endres, M.; Schmitz, D.; Jeworowski, L. M.; Richter, A.; Schmidt, M. L.; Schwarz, T.; Müller, M. A.; Drosten, C.; Wendisch, D.; Sander, L. E.; Osterrieder, N.; Wilson, I. A.; Prüss, H. A Therapeutic Non-Self-Reactive SARS-CoV-2 Antibody Protects from Lung Pathology in a COVID-19 Hamster Model. Cell 2020, 183 (4), 1058–1069.e19.

(39) Hansen, J.; Baum, A.; Pascal, K. E.; Russo, V.; Giordano, S.; Wloga, E.; Fulton, B. O.; Yan, Y.; Koon, K.; Patel, K.; Chung, K. M.; Hermann, A.; Ullman, E.; Cruz, J.; Rafique, A.; Huang, T.; Fairhurst, J.; Libertiny, C.; Malbec, M.; Lee, W. Y.; Welsh, R.; Farr, G.; Pennington, S.; Deshpande, D.; Cheng, J.; Watty, A.; Bouffard, P.; Babb, R.; Levenkova, N.; Chen, C.; Zhang, B.; Hernandez, A. R.; Saotome, K.; Zhou, Y.; Franklin, M.; Sivapalasingam, S.; Lye, D. C.; Weston, S.; Logue, J.; Haupt, R.; Frieman, M.; Chen, G.; Olson, W.; Murphy, A. J.; Stahl, N.; Yancopoulos, G. D.; Kyratsous, C. A. Studies in Humanized Mice and Convalescent Humans Yield a SARS-CoV-2 Antibody Cocktail. Science (80-. ). 2020, 369 (6506), 1010–1014.

(40) Yuan, M.; Liu, H.; Wu, N. C.; Lee, C. C. D.; Zhu, X.; Zhao, F.; Huang, D.; Yu, W.; Hua, Y.; Tien, H.; Rogers, T. F.; Landais, E.; Sok, D.; Jardine, J. G.; Burton, D. R.; Wilson, I. A. Structural Basis of a Shared Antibody Response to SARS-CoV-2. Science (80-. ). 2020, 369 (6507), 1119–1123.

(41) Baral, P.; Bhattarai, N.; Hossen, M. L.; Stebliankin, V.; Gerstman, B. S.; Narasimhan, G.; Chapagain, P. P. Mutation-Induced Changes in the Receptor-Binding Interface of the SARS-CoV-2 Delta Variant B.1.617.2 and Implications for Immune Evasion. Biochem. Biophys. Res. Commun. 2021, 574, 14–19.

(42) Bhattarai, N.; Baral, P.; Gerstman, B. S.; Chapagain, P. P. Structural and Dynamical Differences in the Spike Protein RBD in the SARS-CoV-2 Variants B.1.1.7 and B.1.351. J. Phys. Chem. B 2021, 125 (26), 7101–7107.

(43) Williams, J. K.; Wang, B.; Sam, A.; Hoop, C. L.; Case, D. A.; Baum, J. Molecular Dynamics Analysis of a Flexible Loop at the Binding Interface of the SARS-CoV-2 Spike Protein Receptor-Binding Domain. Proteins Struct. Funct. Bioinforma. 2021.

(44) Nelson, G.; Buzko, O.; Bassett, A.; Spilman, P.; Niazi, K.; Rabizadeh, S.; Soon-Shiong, P. Millisecond-Scale Molecular Dynamics Simulation of Spike RBD Structure Reveals Evolutionary Adaption of SARS-CoV-2 to Stably Bind ACE2. bioRxiv 2020, 2020.12.11.422055.

(45) Tegally, H.; Wilkinson, E.; Giovanetti, M.; Iranzadeh, A.; Fonseca, V.; Giandhari, J.; Doolabh, D.; Pillay, S.; San, E. J.; Msomi, N.; Mlisana, K.; Gottberg, A. von; Walaza, S.; Allam, M.; Ismail, A.; Mohale, T.; Glass, A. J.; Engelbrecht, S.; Zyl, G. Van; Preiser, W.; Petruccione, F.; Sigal, A.; Hardie, D.; Marais, G.; Hsiao, M.; Korsman, S.; Davies, M.-A.; Tyers, L.; Mudau, I.; York, D.; Maslo, C.; Goedhals, D.; Abrahams, S.; Laguda-Akingba, O.; Alisoltani-Dehkordi, A.; Godzik, A.; Wibmer, C. K.; Sewell, B. T.; Lourenço, J.; Alcantara, L. C. J.; Pond, S. L. K.; Weaver, S.; Martin, D.; Lessells, R. J.; Bhiman, J. N.; Williamson, C.; Oliveira, T. de. Emergence and Rapid Spread of a New Severe Acute Respiratory Syndrome-Related Coronavirus 2 (SARS-CoV-2) Lineage with Multiple Spike Mutations in South Africa. medRxiv 2020, 10, 2020.12.21.20248640.

(46) Cherian, S.; Potdar, V.; Jadhav, S.; Yadav, P.; Gupta, N.; Das, M.; Rakshit, P.; Singh, S.; Abraham, P.; Panda, S.; Team, N. SARS-CoV-2 Spike Mutations, L452R, T478K, E484Q and P681R, in the Second Wave of COVID-19 in Maharashtra, India. Microorganisms 2021, 9 (7), 1542.

(47) Science Brief: Emerging SARS-CoV-2 Variants | CDC https://www.cdc.gov/coronavirus/2019-ncov/science/science-briefs/scientific-brief-emerging-variants.html (accessed Sep 30, 2021).

(48) Abdool Karim, S. S.; de Oliveira, T. New SARS-CoV-2 Variants — Clinical, Public Health, and Vaccine Implications. N. Engl. J. Med. 2021, 384 (19), 1866–1868.

(49) Shah, M.; Ahmad, B.; Choi, S.; Woo, H. G. Mutations in the SARS-CoV-2 Spike RBD Are Responsible for Stronger ACE2 Binding and Poor Anti-SARS-CoV MAbs Cross-Neutralization. Comput. Struct. Biotechnol. J. 2020, 18, 3402–3414.

(50) Abraham, M. J.; Murtola, T.; Schulz, R.; Páll, S.; Smith, J. C.; Hess, B.; Lindah, E. Gromacs: High Performance Molecular Simulations through Multi-Level Parallelism from Laptops to Supercomputers. SoftwareX 2015, 1-2, 19–25.

(51) Lindahl; Abraham; Hess; Spoel, van der. GROMACS 2020.3 Source Code. 2020.

(52) Maier, J. A.; Martinez, C.; Kasavajhala, K.; Wickstrom, L.; Hauser, K. E.; Simmerling, C. Ff14SB: Improving the Accuracy of Protein Side Chain and Backbone Parameters from Ff99SB. J. Chem. Theory Comput. 2015, 11 (8), 3696–3713.

(53) Mark, P.; Nilsson, L. Structure and Dynamics of the TIP3P, SPC, and SPC/E Water Models at 298 K. J. Phys. Chem. A 2001, 105 (43), 9954–9960.

(54) Schrödinger, LLC. The {PyMOL} Molecular Graphics System, Version~1.8; 2015.

(55) Berendsen, H. J. C.; Postma, J. P. M.; Van Gunsteren, W. F.; Dinola, A.; Haak, J. R. Molecular Dynamics with Coupling to an External Bath. J. Chem. Phys. 1984, 81 (8), 3684–3690.

(56) Bussi, G.; Donadio, D.; Parrinello, M. Canonical Sampling through Velocity Rescaling. J. Chem. Phys. 2007, 126 (1), 014101.

(57) Parrinello, M.; Rahman, A. Polymorphic Transitions in Single Crystals: A New Molecular Dynamics Method. J. Appl. Phys. 1981, 52 (12), 7182–7190.

(58) Darden, T.; York, D.; Pedersen, L. Particle Mesh Ewald: An N·log(N) Method for Ewald Sums in Large Systems. J. Chem. Phys. 1993, 98 (12), 10089–10092.

(59) Essmann, U.; Perera, L.; Berkowitz, M. L.; Darden, T.; Lee, H.; Pedersen, L. G. A Smooth Particle Mesh Ewald Method. J. Chem. Phys. 1995, 103 (19), 8577–8593.

(60) Hess, B.; Bekker, H.; Berendsen, H. J. C.; Fraaije, J. G. E. M. LINCS: A Linear Constraint Solver for Molecular Simulations. J. Comput. Chem. 1997, 18 (12), 1463–1472.

(61) Humphrey, W.; Dalke, A.; Schulten, K. VMD: Visual Molecular Dynamics. J. Mol. Graph. 1996, 14 (1), 33–38.

(62) Pettersen, E. F.; Goddard, T. D.; Huang, C. C.; Couch, G. S.; Greenblatt, D. M.; Meng, E. C.; Ferrin, T. E. UCSF Chimera?A Visualization System for Exploratory Research and Analysis. J. Comput. Chem. 2004, 25 (13), 1605–1612.

(63) Jolliffe, I. T. Principal Component Analysis; Springer Series in Statistics; Springer-Verlag: New York, 2002.

(64) Jolliffe, I. T.; Cadima, J. Principal Component Analysis: A Review and Recent Developments. Philos. Trans. R. Soc. A Math. Phys. Eng. Sci. 2016, 374 (2065).

(65) Campos, S. R. R.; Baptista, A. M. Conformational Analysis in a Multidimensional Energy Landscape: Study of an Arginylglutamate Repeat. J. Phys. Chem. B 2009, 113 (49), 15989–16001.

(66) Rencher, A. C. Methods of Multivariate Analysis. 2002, 708.

(67) Michaud-Agrawal, N.; Denning, E. J.; Woolf, T. B.; Beckstein, O. MDAnalysis: A Toolkit for the Analysis of Molecular Dynamics Simulations. J. Comput. Chem. 2011, 32 (10), 2319–2327.

(68) Silverman, B. W. Density Estimation for Statistics and Data Analysis; Routledge, 2018.

(69) Campos, S. R. R.; Baptista, A. M. Molecular Simulation Lab — In-House Software https://www.itqb.unl.pt/labs/molecular-simulation/in-house-software (accessed Nov 29, 2021).

(70) Becker, O. M.; Karplus, M. The Topology of Multidimensional Potential Energy Surfaces: Theory and Application to Peptide Structure and Kinetics. J. Chem. Phys. 1998, 106 (4), 1495.

(71) Stillinger, F. H.; Weber, T. A. Packing Structures and Transitions in Liquids and Solids. Science (80-. ). 1984, 225 (4666), 983–989.

(72) Contreras-Riquelme, S.; Garate, J.-A.; Perez-Acle, T.; Martin, A. J. M. RIP-MD: A Tool to Study Residue Interaction Networks in Protein Molecular Dynamics. PeerJ 2018, 6 (12), e5998.

(73) Shannon, P.; Markiel, A.; Ozier, O.; Baliga, N. S.; Wang, J. T.; Ramage, D.; Amin, N.; Schwikowski, B.; Ideker, T. Cytoscape: A Software Environment for Integrated Models of Biomolecular Interaction Networks. Genome Res. 2003, 13 (11), 2498.

(74) Wu, N. C.; Yuan, M.; Liu, H.; Lee, C.-C. D.; Zhu, X.; Bangaru, S.; Torres, J. L.; Caniels, T. G.; Brouwer, P. J. M.; Gils, M. J. van; Sanders, R. W.; Ward, A. B.; Wilson, I. A. An Alternative Binding Mode of IGHV3-53 Antibodies to the SARS-CoV-2 Receptor Binding Domain. Cell Rep. 2020, 33 (3).

(75) McCallum, M.; Walls, A. C.; Sprouse, K. R.; Bowen, J. E.; Rosen, L.; Dang, H. V.; deMarco, A.; Franko, N.; Tilles, S. W.; Logue, J.; Miranda, M. C.; Ahlrichs, M.; Carter, L.; Snell, G.; Pizzuto, M. S.; Chu, H. Y.; Voorhis, W. C. Van; Corti, D.; Veesler, D. Molecular Basis of Immune Evasion by the Delta and Kappa SARS-CoV-2 Variants. bioRxiv 2021, 2021.08.11.455956.

(76) Tian, F.; Tong, B.; Sun, L.; Shi, S.; Zheng, B.; Wang, Z.; Dong, X.; Zheng, P. N501Y Mutation of Spike Protein in SARS-CoV-2 Strengthens Its Binding to Receptor ACE2. Elife 2021, 10.

(77) Laffeber, C.; de Koning, K.; Kanaar, R.; Lebbink, J. H. G. Experimental Evidence for Enhanced Receptor Binding by Rapidly Spreading SARS-CoV-2 Variants. J. Mol. Biol. 2021, 433 (15), 167058.

(78) Supasa, P.; Zhou, D.; Dejnirattisai, W.; Liu, C.; Mentzer, A. J.; Ginn, H. M.; Zhao, Y.; Duyvesteyn, H. M. E.; Nutalai, R.; Tuekprakhon, A.; Wang, B.; Paesen, G. C.; Slon-Campos, J.; López-Camacho, C.; Hallis, B.; Coombes, N.; Bewley, K. R.; Charlton, S.; Walter, T. S.; Barnes, E.; Dunachie, S. J.; Skelly, D.; Lumley, S. F.; Baker, N.; Shaik, I.; Humphries, H. E.; Godwin, K.; Gent, N.; Sienkiewicz, A.; Dold, C.; Levin, R.; Dong, T.; Pollard, A. J.; Knight, J. C.; Klenerman, P.; Crook, D.; Lambe, T.; Clutterbuck, E.; Bibi, S.; Flaxman, A.; Bittaye, M.; Belij-Rammerstorfer, S.; Gilbert, S.; Hall, D. R.; Williams, M. A.; Paterson, N. G.; James, W.; Carroll, M. W.; Fry, E. E.; Mongkolsapaya, J.; Ren, J.; Stuart, D. I.; Screaton, G. R. Reduced Neutralization of SARS-CoV-2 B.1.1.7 Variant by Convalescent and Vaccine Sera. Cell 2021, 184 (8), 2201–2211.e7.

(79) Wirnsberger, G.; Monteil, V.; Eaton, B.; Postnikova, E.; Murphy, M.; Braunsfeld, B.; Crozier, I.; Kricek, F.; Niederhöfer, J.; Schwarzböck, A.; Breid, H.; Jimenez, A. S.; Bugajska-Schretter, A.; Dohnal, A.; Ruf, C.; Gugenberger, R.; Hagelkruys, A.; Montserrat, N.; Holbrook, M. R.; Oostenbrink, C.; Shoemaker, R. H.; Mirazimi, A.; Penninger, J. M. Clinical Grade ACE2 as a Universal Agent to Block SARS-CoV-2 Variants. bioRxiv 2021, 2021.09.10.459744.

(80) Souza, A. S. de; Amorim, V. M. de F.; Guardia, G. D. A.; Santos, F. R. C. dos; Santos, F. F. dos; Souza, R. F. de; Juvenal, G. de A.; Huang, Y.; Ge, P.; Jiang, Y.; Paudel, P.; Ulrich, H.; Galante, P. A. F.; Guzzo, C. R. Molecular Dynamics Analysis of Fast-Spreading Severe Acute Respiratory Syndrome Coronavirus 2 Variants and Their Effects in the Interaction with Human Angiotensin-Converting Enzyme 2. bioRxiv 2021, 2021.06.14.448436.

(81) Saville, J. W.; Mannar, D.; Zhu, X.; Srivastava, S. S.; Berezuk, A. M.; Demers, J.-P.; Zhou, S.; Tuttle, K. S.; Sekirov, I.; Kim, A.; Li, W.; Dimitrov, D. S.; Subramaniam, S. Structural and Biochemical Rationale for Enhanced Spike Protein Fitness in Delta and Kappa SARS-CoV-2 Variants. bioRxiv 2021, 2021.09.02.458774.

(82) Socher, E.; Conrad, M.; Heger, L.; Paulsen, F.; Sticht, H.; Zunke, F.; Arnold, P. Computational Decomposition Reveals Reshaping of the SARS-CoV-2–ACE2 Interface among Viral Variants Expressing the N501Y Mutation. J. Cell. Biochem. 2021.

(83) Ali, F.; Kasry, A.; Amin, M. The New SARS-CoV-2 Strain Shows a Stronger Binding Affinity to ACE2 Due to N501Y Mutant. Med. Drug Discov. 2021, 10, 100086.

(84) Luan, B.; Wang, H.; Huynh, T. Enhanced Binding of the N501Y-Mutated SARS-CoV-2 Spike Protein to the Human ACE2 Receptor: Insights from Molecular Dynamics Simulations. FEBS Lett. 2021, 595 (10), 1454–1461.

(85) Ahmed, W.; Philip, A. M.; Biswas, K. H. Stable Interaction Of The UK B.1.1.7 Lineage SARS-CoV-2 S1 Spike N501Y Mutant With ACE2 Revealed By Molecular Dynamics Simulation. bioRxiv 2021, 2021.01.07.425307.

(86) Nelson, G.; Buzko, O.; Spilman, P.; Niazi, K.; Rabizadeh, S.; Soon-Shiong, P. Molecular Dynamic Simulation Reveals E484K Mutation Enhances Spike RBD-ACE2 Affinity and the Combination of E484K, K417N and N501Y Mutations (501Y.V2 Variant) Induces Conformational Change Greater than N501Y Mutant Alone, Potentially Resulting in an Escape Mutant. bioRxiv 2021, 2021.01.13.426558.

(87) CoVariants https://covariants.org/variants/21K.Omicron (accessed Nov 29, 2021).

(88) Walls, A. C.; Park, Y. J.; Tortorici, M. A.; Wall, A.; McGuire, A. T.; Veesler, D. Structure, Function, and Antigenicity of the SARS-CoV-2 Spike Glycoprotein. Cell 2020, 181 (2), 281–292.e6.

(89) Norman, A.; Franck, C.; Christie, M.; Hawkins, P. M. E.; Patel, K.; Ashhurst, A. S.; Aggarwal, A.; Low, J. K. K.; Siddiquee, R.; Ashley, C. L.; Steain, M.; Triccas, J. A.; Turville, S.; Mackay, J. P.; Passioura, T.; Payne, R. J. Discovery of Cyclic Peptide Ligands to the SARS-CoV-2 Spike Protein Using MRNA Display. ACS Cent. Sci. 2021, 7 (6), 1001–1008.

(90) Jiang, W.; Wang, J.; Jiao, S.; Gu, C.; Xu, W.; Chen, B.; Wang, R.; Chen, H.; Xie, Y.; Wang, A.; Li, G.; Zeng, D.; Zhang, J.; Zhang, M.; Wang, S.; Wang, M.; Gui, X. Characterization of MW06, a Human Monoclonal Antibody with Cross-Neutralization Activity against Both SARS-CoV-2 and SARS-CoV. MAbs 2021, 13 (1), 1953683.

(91) Yao, H.; Sun, Y.; Deng, Y.-Q.; Wang, N.; Tan, Y.; Zhang, N.-N.; Li, X.-F.; Kong, C.; Xu, Y.-P.; Chen, Q.; Cao, T.-S.; Zhao, H.; Yan, X.; Cao, L.; Lv, Z.; Zhu, D.; Feng, R.; Wu, N.; Zhang, W.; Hu, Y.; Chen, K.; Zhang, R.-R.; Lv, Q.; Sun, S.; Zhou, Y.; Yan, R.; Yang, G.; Sun, X.; Liu, C.; Lu, X.; Cheng, L.; Qiu, H.; Huang, X.-Y.; Weng, T.; Shi, D.; Jiang, W.; Shao, J.; Wang, L.; Zhang, J.; Jiang, T.; Lang, G.; Qin, C.-F.; Li, L.; Wang, X. Rational Development of a Human Antibody Cocktail That Deploys Multiple Functions to Confer Pan-SARS-CoVs Protection. Cell Res. 2020 311 2020, 31 (1), 25–36.

(92) Fu, D.; Zhang, G.; Wang, Y.; Zhang, Z.; Hu, H.; Shen, S.; Wu, J.; Li, B.; Li, X.; Fang, Y.; Liu, J.; Wang, Q.; Zhou, Y.; Wang, W.; Li, Y.; Lu, Z.; Wang, X.; Nie, C.; Tian, Y.; Chen, D.; Wang, Y.; Zhou, X.; Wang, Q.; Yu, F.; Zhang, C.; Deng, C.; Zhou, L.; Guan, G.; Shao, N.; Lou, Z.; Deng, F.; Zhang, H.; Chen, X.; Wang, M.; Liu, L.; Rao, Z.; Guo, Y. Structural Basis for SARS-CoV-2 Neutralizing Antibodies with Novel Binding Epitopes. PLOS Biol. 2021, 19 (5), e3001209.

