## SupplementalMaterial for "SARS-CoV-2 variants impact RBD conformational dynamics and ACE2 accessibility"

**Videos S1 to S4. RBD and RBM conformation dynamics for the wt, alpha, beta, and delta variants.** Trajectory samples recovered from the AA MD simulations of the *wt*, alpha, beta and delta RBD in water. The ridge region of the RBD is colored in red. Residues of interest are labelled at the start of the video. Renderization done with VMD, with the positions averaged over 5 consecutive frames.

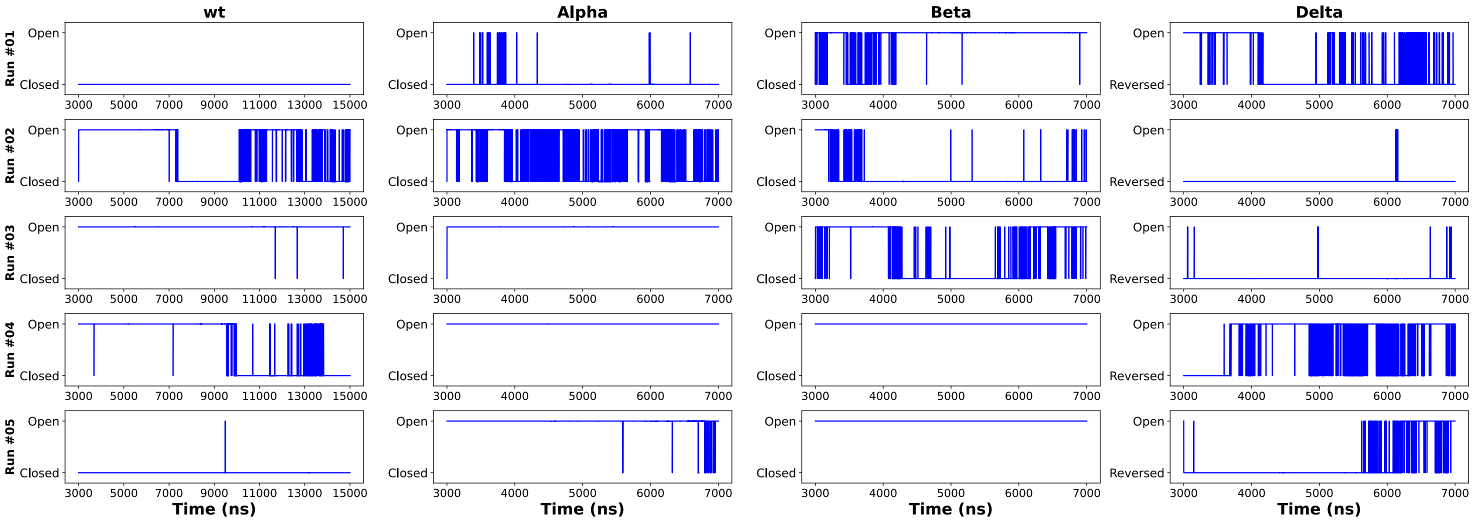


**Figure S1. RBD open – closed dynamics over simulation time.** RBD open – closed dynamics as determined by analysis of the conformational basins (Supplementary Table S1) recovered by PCA. Data shown for the five replicas for each variant tested. The first 3 µs of simulation were used for equilibration.

**Table S1. Energy surface landscape analysis from 2D PCA of SARS-CoV-2 RBD conformational dynamics in water.** Energy surface landscape analysis and defined basins for each of the tested RBD variants. Energy minima, frame percentage and loop conformation for each of the basins is also given. Overall analysis of "open" vs. "closed" conformation is also shown. 95 % Confidence intervals (CI) were calculated with bootstrap resampling from the frame percentages recovered from the individual simulation replicas. Representative structures can be seen in figures S8, S9, S10 and S11.

| | Basin | Free Energy | $\langle E\rangle/k_BT$ | S/R | $E_{min}/k_BT$ | Frame percentage (%) | Loop Conformation | Time in "closed" state (%) ± CI (95%) | Time in "open" state (%) ± CI (95%) | "closing" ΔΔG (kJ/mol) |
|---|---|---|---|---|---|---|---|---|---|---|
| | 0 | -5.16 | 1.36 | 6.52 | 0.44 | 37.69 | "Closed" | | | |
| | 1 | -4.67 | 0.99 | 5.67 | 0.26 | 23.01 | "Open" | | | |
| | 2 | -4.41 | 0.57 | 4.98 | 0.00 | 17.66 | "Closed" | | | |
| | 3 | -4.20 | 1.04 | 5.24 | 0.44 | 14.42 | "Open" | | | |
| | 4 | -3.28 | 3.62 | 6.90 | 2.86 | 5.77 | "Open" | | | |
| ***WT*** | 5 | -1.57 | 4.07 | 5.64 | 3.61 | 1.04 | "Open" | $55.50^{+2.12}_{-2.93}$ | $44.49^{+0.93}_{-1.12}$ | $0.5^{+0.21}_{-0.13}$ |
| | 6 | 0.79 | 5.30 | 4.51 | 5.02 | 0.13 | "Open" | | | |
| | 7 | 0.83 | 5.42 | 4.60 | 5.21 | 0.14 | "Closed" | | | |
| | 8 | 0.91 | 5.39 | 4.48 | 5.01 | 0.12 | "Open" | | | |
| | 9 | 2.70 | 5.06 | 2.37 | 5.01 | 0.01 | "Closed" | | | |
| | | | | | **Total** | 99.99 | | | | |
| | 0 | -5.20 | 1.02 | 6.22 | 0.00 | 39.18 | "Open" | | | |
| | 1 | -4.64 | 1.47 | 6.11 | 0.35 | 22.41 | "Open" | | | |
| | 2 | -4.37 | 1.84 | 6.21 | 0.83 | 17.23 | "Closed" | | | |
| | 3 | -3.82 | 0.96 | 4.78 | 0.35 | 9.85 | "Closed" | | | |
| | 4 | -3.10 | 2.77 | 5.86 | 1.83 | 4.82 | "Open" | | | |
| **Alpha (N501Y)** | 5 | -2.71 | 3.88 | 6.58 | 3.01 | 3.28 | "Open" | $27.31^{+1.79}_{-2.04}$ | $72.64^{+1.16}_{-1.43}$ | $-2.44^{+0.21}_{-0.23}$ |
| | 6 | -2.34 | 4.10 | 6.44 | 3.11 | 2.41 | "Open" | | | |
| | 7 | -0.28 | 5.21 | 5.49 | 4.93 | 0.29 | "Open" | | | |
| | 8 | -0.11 | 4.43 | 4.54 | 4.05 | 0.25 | "Open" | | | |
| | 9 | 0.11 | 5.46 | 5.35 | 5.10 | 0.23 | "Closed" | | | |
| | | | | | **Total** | 99.95 | | | | |
| | 0 | -4.95 | 0.58 | 5.53 | 0.00 | 23.73 | "Open" | | | |
| | 1 | -4.70 | 0.46 | 5.16 | 0.09 | 18.41 | "Open" | | | |
| | 2 | -4.38 | 1.31 | 5.69 | 0.90 | 13.43 | "Closed" | | | |
| | 3 | -3.98 | 1.32 | 5.30 | 0.90 | 9.03 | "Closed" | | | |
| | 4 | -3.83 | 2.70 | 6.53 | 1.93 | 7.72 | "Closed" | | | |
| | 5 | -3.78 | 2.86 | 6.64 | 2.02 | 7.36 | "Open" | | | |
| | 6 | -3.36 | 2.93 | 6.30 | 2.30 | 4.86 | "Open" | | | |
| **Beta (K417N E484K N501Y)** | 7 | -3.20 | 2.20 | 5.40 | 1.68 | 4.14 | "Open" | $30.18^{+1.24}_{-1.74}$ | $69.81^{+0.66}_{-0.73}$ | $-2.09^{+0.13}_{-0.15}$ |
| | 8 | -2.93 | 4.08 | 7.02 | 3.32 | 3.21 | "Open" | | | |
| | 9 | -2.71 | 3.66 | 6.37 | 3.11 | 2.54 | "Open" | | | |
| | 10 | -2.58 | 3.53 | 6.12 | 2.98 | 2.23 | "Open" | | | |
| | 11 | -2.14 | 3.70 | 5.85 | 3.28 | 1.44 | "Open" | | | |
| | 12 | -1.74 | 3.78 | 5.52 | 3.48 | 0.95 | "Open" | | | |
| | 13 | -1.72 | 3.99 | 5.71 | 3.70 | 0.94 | "Open" | | | |
| | | | | | **Total** | 99.99 | | | | |
| | 0 | -5.74 | 1.03 | 6.77 | 0.00 | 53.51 | "Open" | | | |
| | 1 | -4.97 | 2.32 | 7.29 | 1.46 | 24.76 | 'Reverse Open" | | | |
| | 2 | -4.13 | 1.70 | 5.84 | 1.20 | 10.75 | 'Reverse Open" | | | |
| **Delta (L452R T478K)** | 3 | -3.46 | 3.17 | 6.63 | 2.62 | 5.50 | "Open" | N/A | 100.00 | N/A |
| | 4 | -2.93 | 3.34 | 6.27 | 2.63 | 3.22 | "Open" | | | |
| | 5 | -1.88 | 4.15 | 6.03 | 3.77 | 1.12 | "Open" | | | |
| | 6 | -1.85 | 4.12 | 5.97 | 3.60 | 1.13 | "Open" | | | |
| | | | | | **Total** | 99.99 | | | | |

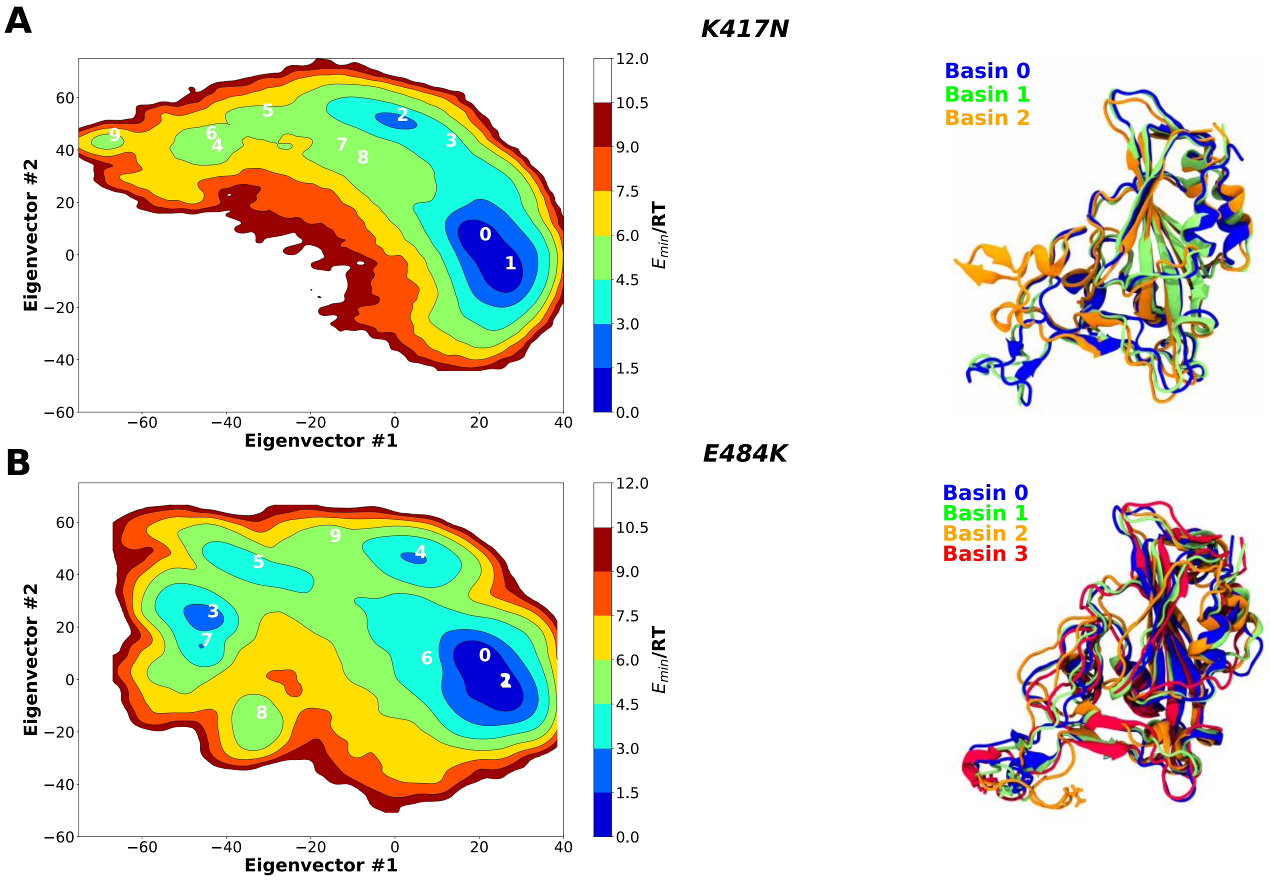


**Figure S2. Two-dimension principal component analysis (PCA) of SARS-CoV-2 RBD's mutants conformational dynamics in water.** Plots of the first two principal components determined from the Cα backbone of the (A) K417N and (B) E484K RBD mutants. Snapshots of the lowest energy structures for selected basins are also shown.

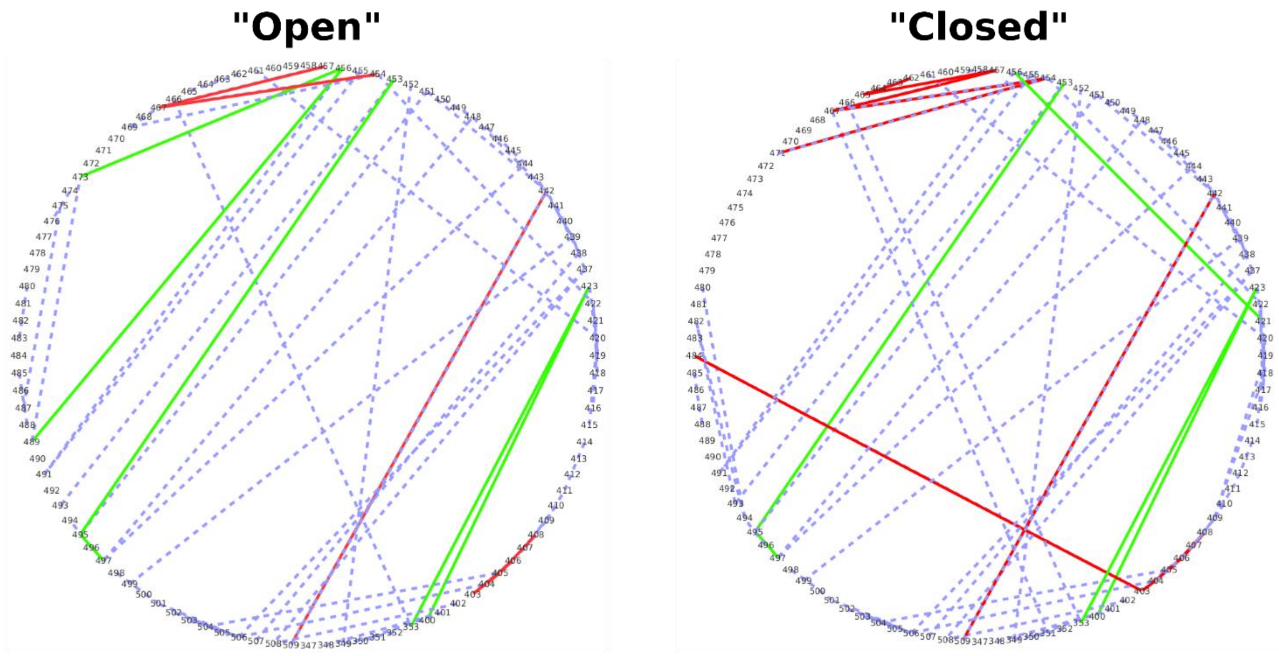


**Figure S3. Residue interaction networks (RINs) for the "open" and "closed" conformations of *wt* RBD.** RINs determined using RIP-MD for the 5000 lowest energy conformations obtained for the most populated "open" and "closed" basins of *wt* RBD. Hydrogen bonds, salt bridges and pi-pi interactions are shown in blue, red and green, respectively.

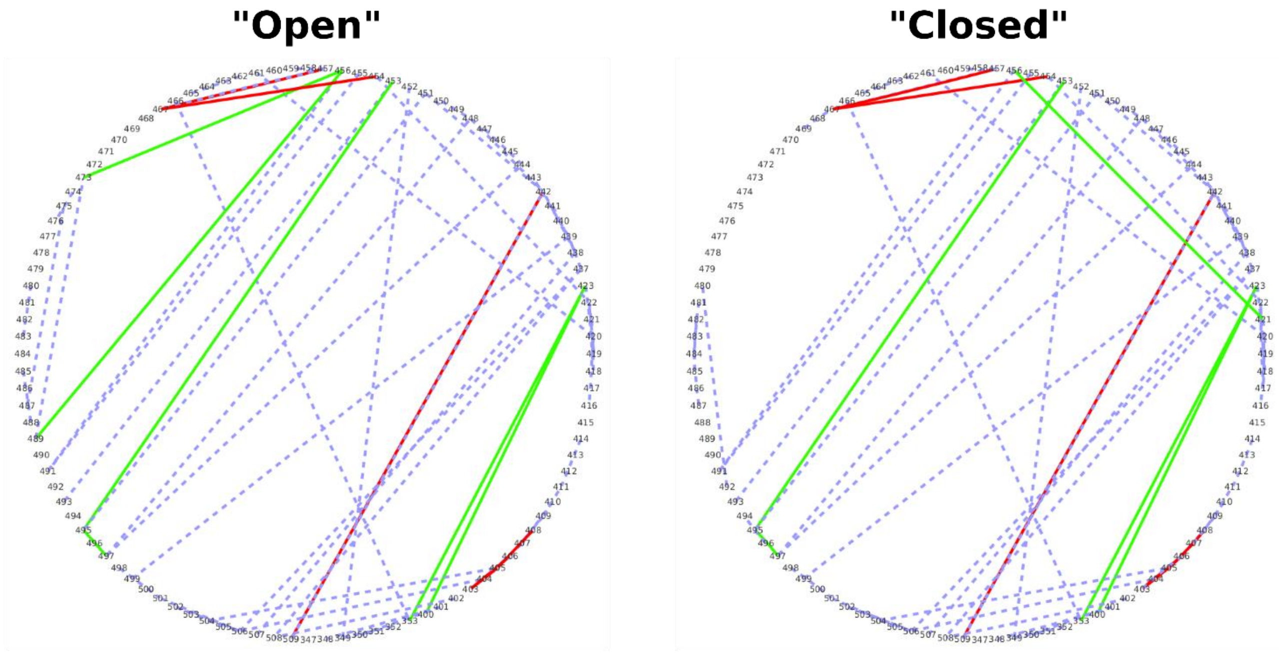


**Figure S4. Residue interaction networks (RINs) for the "open" and "closed" conformations of alpha variant.** RINs determined using RIP-MD for the 5000 lowest energy conformations obtained for the most populated "open" and "closed" basins of alpha RBD. Hydrogen bonds, salt bridges and pi-pi interactions are shown in blue, red and green, respectively.

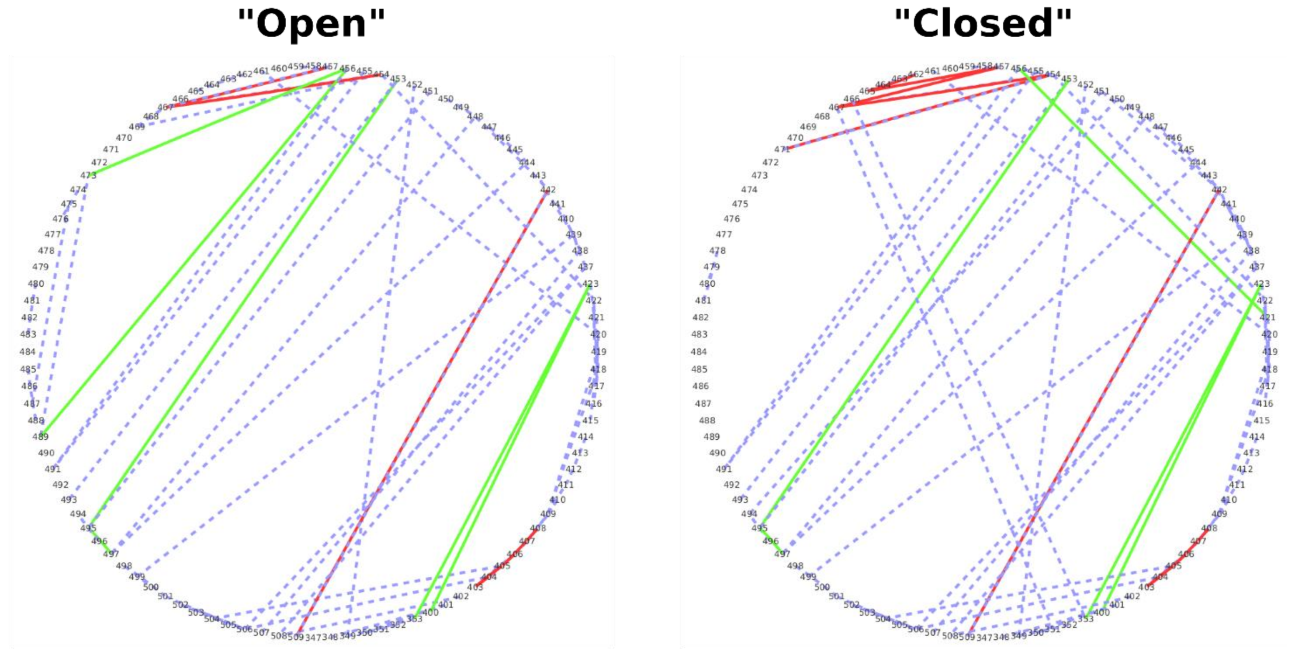


**Figure S5. Residue interaction networks (RINs) for the "open" and "closed" conformations of beta variant.** RINs determined using RIP-MD for the 5000 lowest energy conformations obtained for the most populated "open" and "closed" basins of beta RBD. Hydrogen bonds, salt bridges and pi-pi interactions are shown in blue, red and green, respectively.

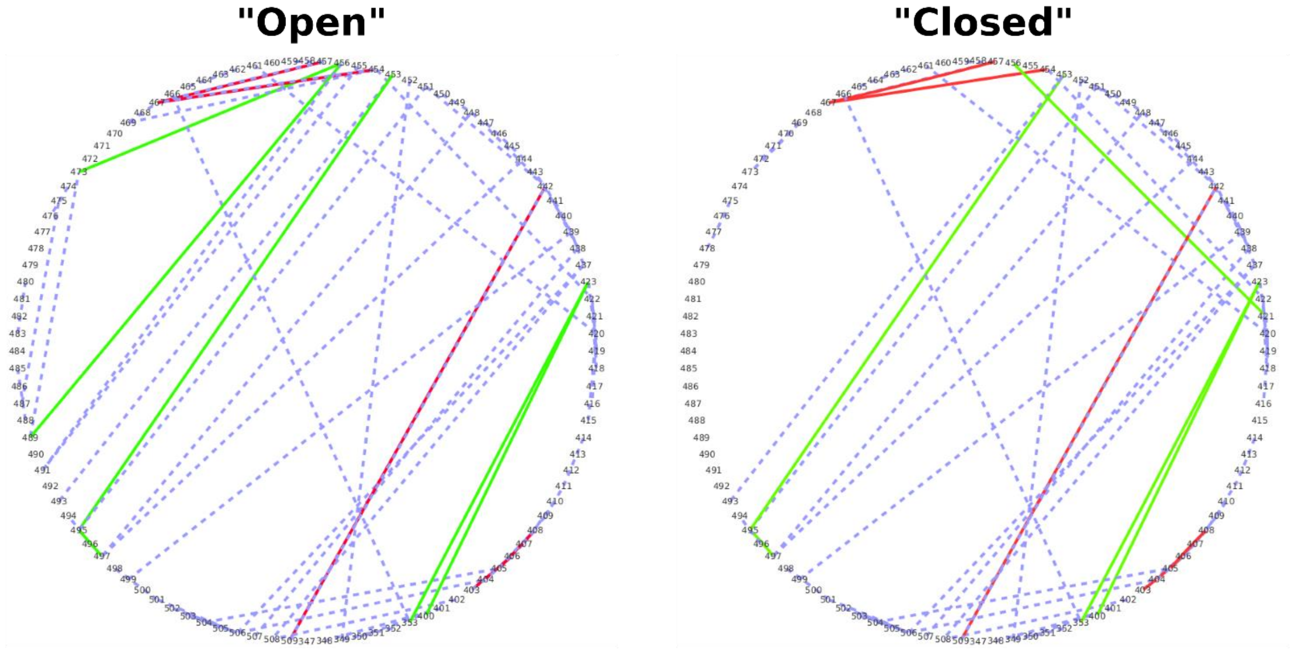


**Figure S6. Residue interaction networks (RINs) for the "open" and "reversed" conformations of delta variant.** RINs determined using RIP-MD for the 5000 lowest energy conformations obtained for "open" and "reversed" basins of alpha RBD. Hydrogen bonds, salt bridges and pi-pi interactions are shown in blue, red and green, respectively.

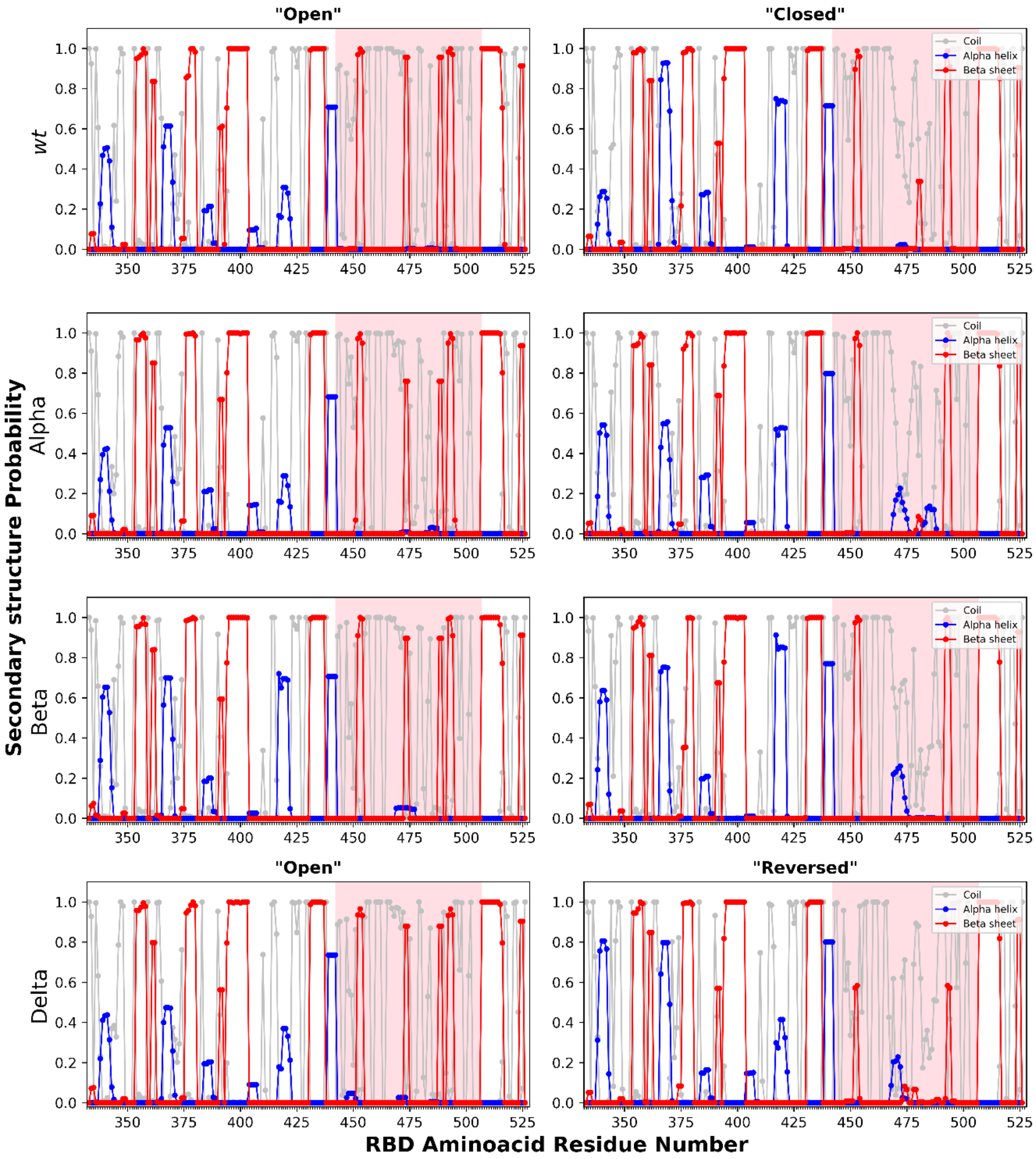


**Figure S7. Secondary structure of *wt*, alpha, beta and delta SARS-CoV-2 RBD simulated in water.** Probability of coil, α-helix and β-sheet secondary structures was obtained using the GROMACS tool gmx do_dssp[1] for both conformations ("open"/"closed " and "open"/"reversed") of all four variants. RBM region is highlighted in pink.

**Table S2. Surface Accessible Surface Area (SASA) analysis of SARS-CoV-2 RBD in water.** SASA values were calculated using the GROMACS tool gmx_sasa[1] for the whole trajectory (Entire trajectory) and for the two major conformations ("open"/"closed " and "open"/"reversed"). Results were also divided in the contribution of hydrophobic atoms, which are the ones with charges [-0.2, 0.2], and hydrophilic, those outside of this range. 95 % Confidence intervals (CI) were calculated with bootstrap resampling.

| Variant | Protein SASA ($nm^2$) | Hydrophobic atoms SASA ($nm^2$) | Hydrophilic atoms SASA ($nm^2$) |
|---|---|---|---|
| | Average ± CI (95%) | Average ± CI (95%) | Average ± CI (95%) |
| ***WT*** | | | |
| *Average* | $107.72^{+0.07}_{-0.07}$ | $55.07^{+0.05}_{-0.05}$ | $52.62^{+0.05}_{-0.05}$ |
| *"Open"* | $109.08^{+0.05}_{-0.05}$ | $55.11^{+0.04}_{-0.04}$ | $53.97^{+0.04}_{-0.05}$ |
| *"Closed"* | $105.52^{+0.05}_{-0.05}$ | $53.68^{+0.04}_{-0.04}$ | $51.57^{+0.04}_{-0.04}$ |
| **Alpha** | | | |
| *Average* | $108.16^{+0.04}_{-0.04}$ | $55.13^{+0.03}_{-0.03}$ | $53.04^{+0.03}_{-0.03}$ |
| *"Open"* | $108.62^{+0.06}_{-0.06}$ | $55.26^{+0.04}_{-0.04}$ | $53.36^{+0.04}_{-0.04}$ |
| *"Closed"* | $102.99^{+0.07}_{-0.07}$ | $52.69^{+0.04}_{-0.04}$ | $50.31^{+0.05}_{-0.05}$ |
| **Beta** | | | |
| *Average* | $108.10^{+0.01}_{-0.01}$ | $54.95^{+0.01}_{-0.01}$ | $53.15^{+0.01}_{-0.01}$ |
| *"Open"* | $108.58^{+0.06}_{-0.06}$ | $53.70^{+0.04}_{-0.04}$ | $53.88^{+0.04}_{-0.04}$ |
| *"Closed"* | $107.65^{+0.08}_{-0.08}$ | $56.32^{+0.05}_{-0.05}$ | $51.33^{+0.05}_{-0.06}$ |
| **Delta** | | | |
| *Average* | $109.62^{+0.11}_{-0.11}$ | $55.46^{+0.08}_{-0.08}$ | $54.16^{+0.08}_{-0.07}$ |
| *"Open"* | $109.58^{+0.05}_{-0.05}$ | $54.87^{+0.04}_{-0.04}$ | $54.70^{+0.04}_{-0.04}$ |
| *"Reversed"* | $106.08^{+0.07}_{-0.07}$ | $53.65^{+0.04}_{-0.04}$ | $52.43^{+0.04}_{-0.04}$ |

**Table S3. Compilation of ACE2-RBD binding kinetics data from recent studies.** Kinetic parameters of ACE2 binding to *wt*, alpha, beta and delta RBD/Spike variants data obtained from SPR and BLI[2–8] .

| | | | *WT* | | | Alpha | | | Beta | | | Delta | | |
|---|---|---|---|---|---|---|---|---|---|---|---|---|---|---|
| Reference | RBD/Spike | Technique | $K_{on}$ ($M^{-1}$ $s^{-1}$) | $K_{off}$ ($s^{-1}$) | $K_d$ (nM) | $K_{on}$ ($M^{-1}$ $s^{-1}$) | $K_{off}$ ($s^{-1}$) | $K_d$ (nM) | $K_{on}$ ($M^{-1}$ $s^{-1}$) | $K_{off}$ ($s^{-1}$) | $K_d$ (nM) | $K_{on}$ ($M^{-1}$ $s^{-1}$) | $K_{off}$ ($s^{-1}$) | $K_d$ (nM) |
| McCallum et al. 2021 [2] | RBD | SPR | $7.70 \times 10^4$ | $6.70 \times 10^{-3}$ | 78 | $7.50 \times 10^4$ | $1.20 \times 10^{-3}$ | 15 | - | - | - | $5.90 \times 10^4$ | $4.30 \times 10^{-3}$ | 63 |
| Tian et al. 2021 [3] | RBD | SPR | $2.50 \times 10^4$ | $2.10 \times 10^{-4}$ | 8.3 | $7.50 \times 10^4$ | $0.37 \times 10^{-4}$ | 0.5 | $6.40 \times 10^4$ | $0.30 \times 10^{-4}$ | 0.5 | - | - | - |
| Laffeber et al. 2021 [4] | RBD | SPR | $4.50 \times 10^5$ | $7.80 \times 10^{-3}$ | 17 | $5.70 \times 10^5$ | $1.30 \times 10^{-3}$ | 2.4 | $7.60 \times 10^5$ | $4.30 \times 10^{-3}$ | 5.8 | - | - | - |
| Supasa et al. 2021 [5] | RBD | SPR | $3.88 \times 10^4$ | $2.92 \times 10^{-3}$ | 75.1 | $6.38 \times 10^4$ | $6.85 \times 10^{-4}$ | 10.7 | - | - | - | - | - | - |
| Wirnsberger et al. 2021 [6] | RBD | SPR | $6.86 \times 10^5$ | $11.0 \times 10^{-3}$ | 16.2 | $4.36 \times 10^5$ | $1.59 \times 10^{-3}$ | 3.72 | $6.86 \times 10^5$ | $4.42 \times 10^{-3}$ | 6.5 | $10.1 \times 10^5$ | $8.23 \times 10^{-3}$ | 8.61 |
| de Souza et al. 2021 [7] | RBD | SPR | $0.90 \times 10^6$ | $91.6 \times 10^{-4}$ | 10.3 | $1.30 \times 10^6$ | $15.5 \times 10^{-4}$ | 1.2 | $1.20 \times 10^6$ | $39.4 \times 10^{-4}$ | 3.3 | | | |
| de Souza et al. 2021 [7] | Spike | SPR | $0.09 \times 10^6$ | $5.80 \times 10^{-4}$ | 6.4 | $0.10 \times 10^6$ | $1.70 \times 10^{-4}$ | 0.1 | $0.30 \times 10^6$ | $3.00 \times 10^{-4}$ | 0.3 | | | |
| Saville et al. 2021 [8] | Spike | BLI | $1.40 \times 10^5$ | $7.09 \times 10^{-4}$ | 5.06 | - | - | - | - | - | - | $1.51 \times 10^5$ | $4.01 \times 10^{-4}$ | 2.65 |
| Yang et al. 2021 [9] | Spike | BLI | $3.43 \times 10^4$ | $10.6 \times 10^{-5}$ | 3.1 | $3.86 \times 10^4$ | $5.25 \times 10^{-5}$ | 1.36 | $7.32 \times 10^4$ | $1.79 \times 10^{-5}$ | 0.25 | $4.39 \times 10^4$ | $1.71 \times 10^{-5}$ | 0.39 |

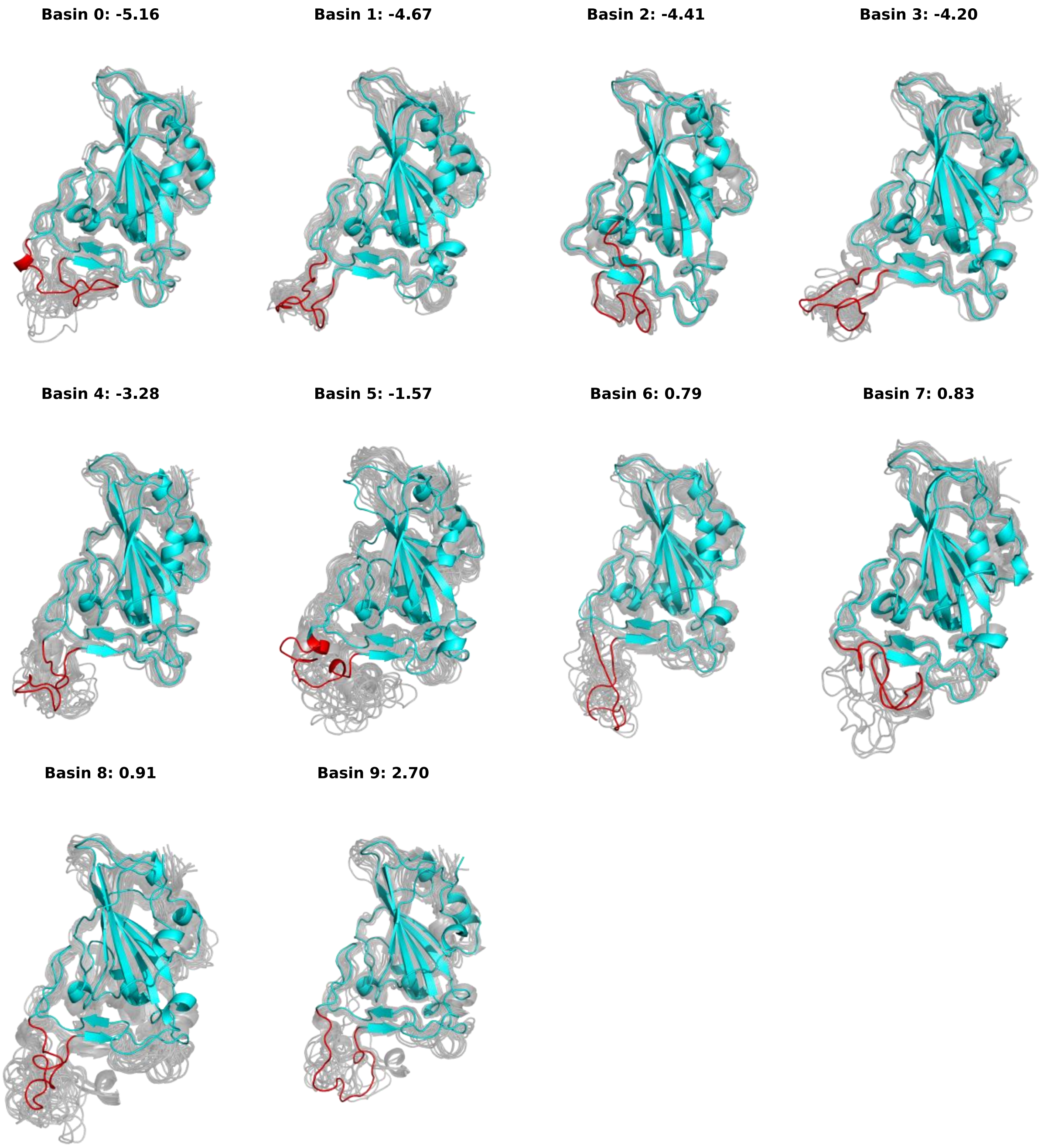


**Figure S8. Snapshots representative of all *wt* RBD PCA basins.** The structures corresponding to the free energy minima of all conformational basins are represented in blue, with the ridge region highlighted in red, together with structures sampled from the same basin (background, gray colored).

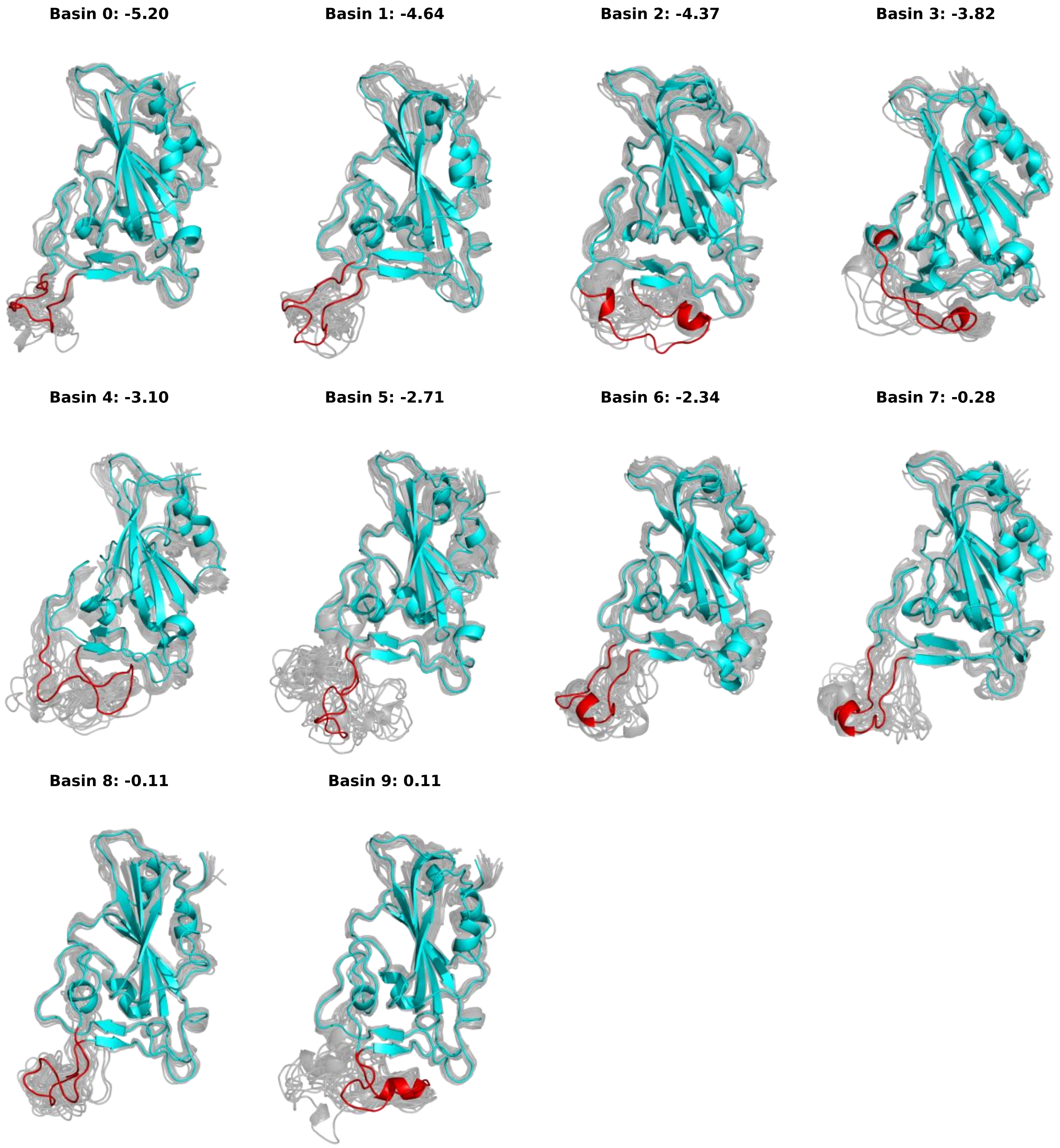


**Figure S9. Structures representative of all alpha RBD PCA basins.** The structures corresponding to the free energy minima of all conformational basins are represented in blue, with the ridge region highlighted in red, together with structures sampled from the same basin (background, gray colored).

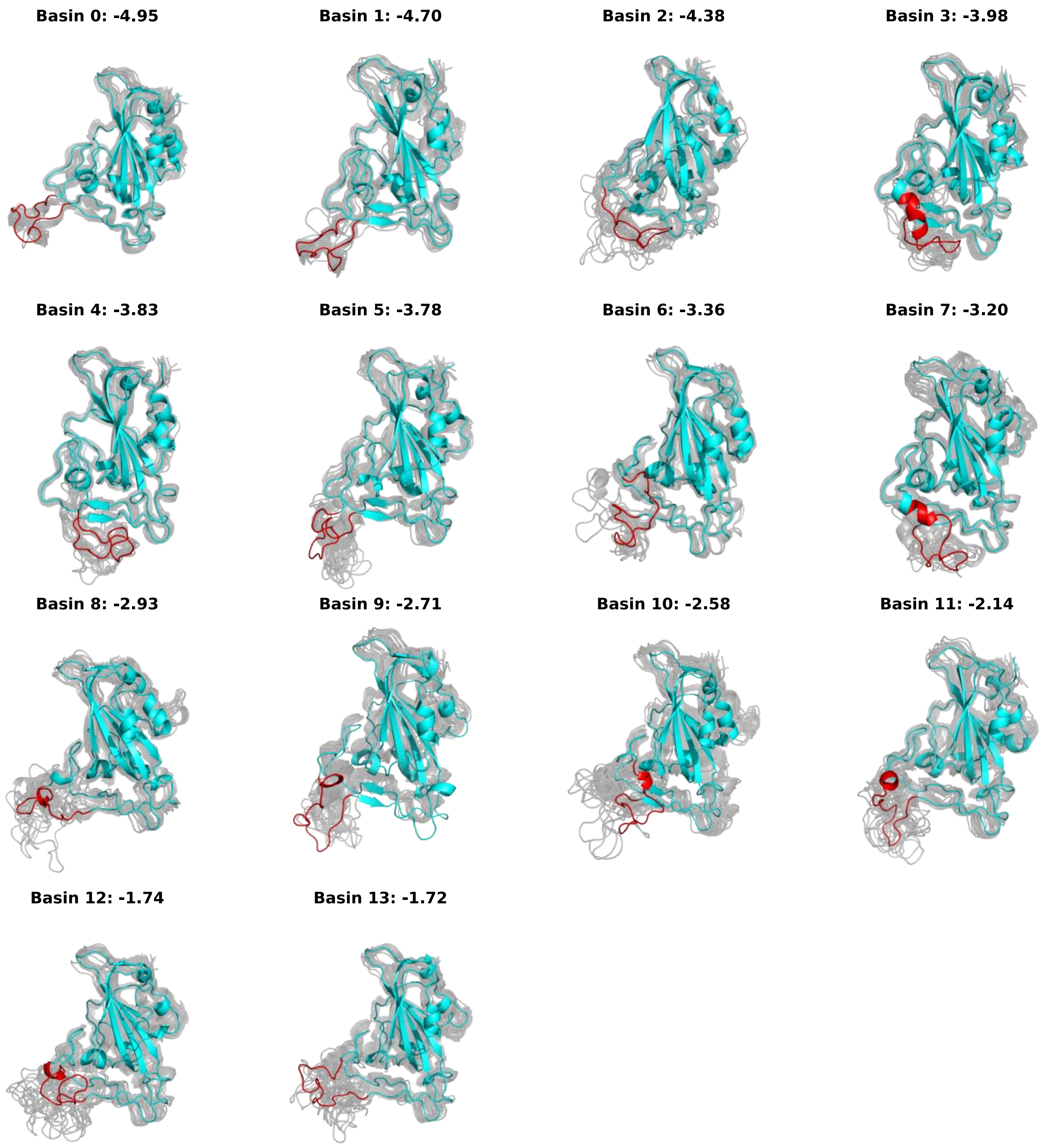


**Figure S10. Structures representative of all beta RBD PCA basins.** The structures corresponding to the free energy minima of all conformational basins are represented in blue, with the ridge region highlighted in red, together with structures sampled from the same basin (background, gray colored).

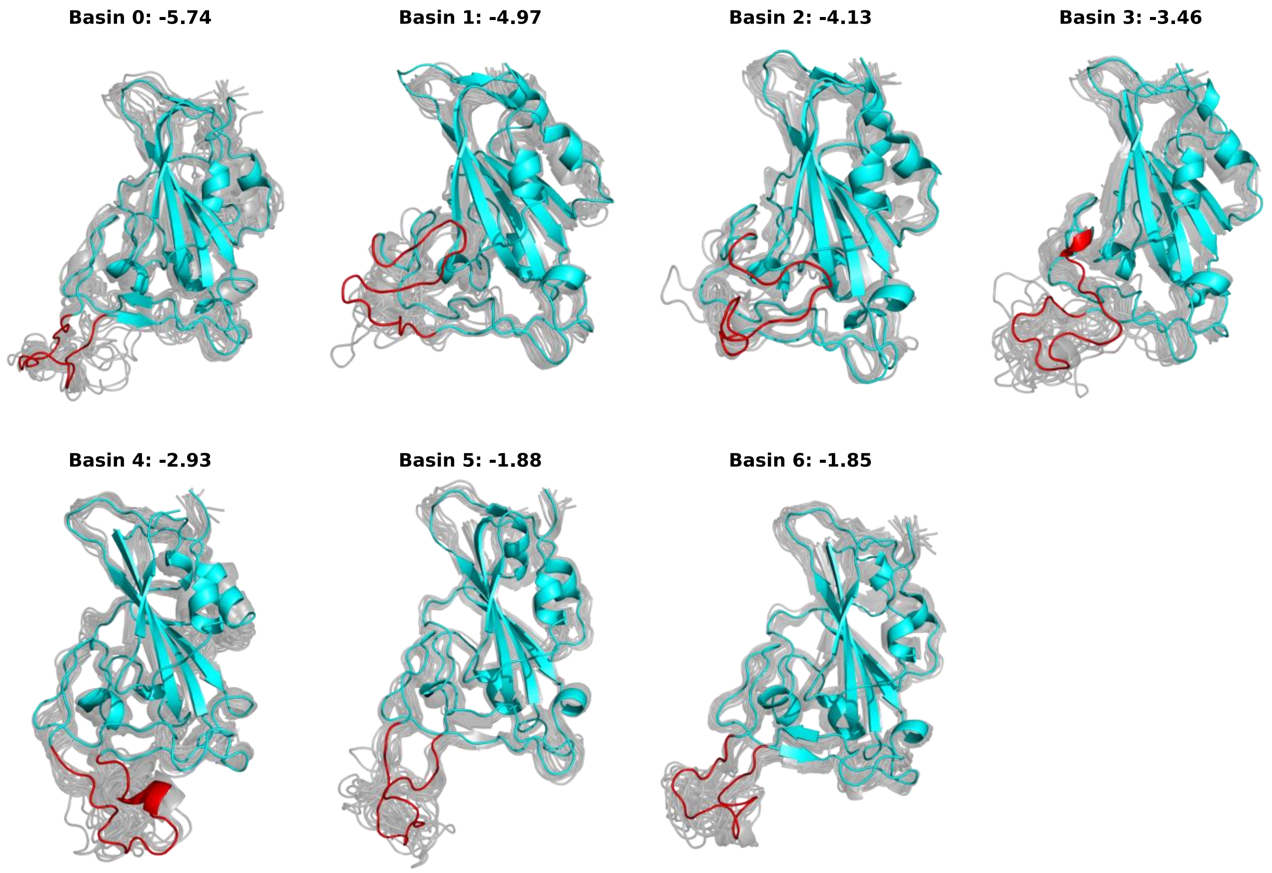


**Figure S11. Structures representative of all delta RBD PCA basins.** The structures corresponding to the free energy minima of all conformational basins are represented in blue, with the ridge region highlighted in red, together with structures sampled from the same basin (background, gray colored).

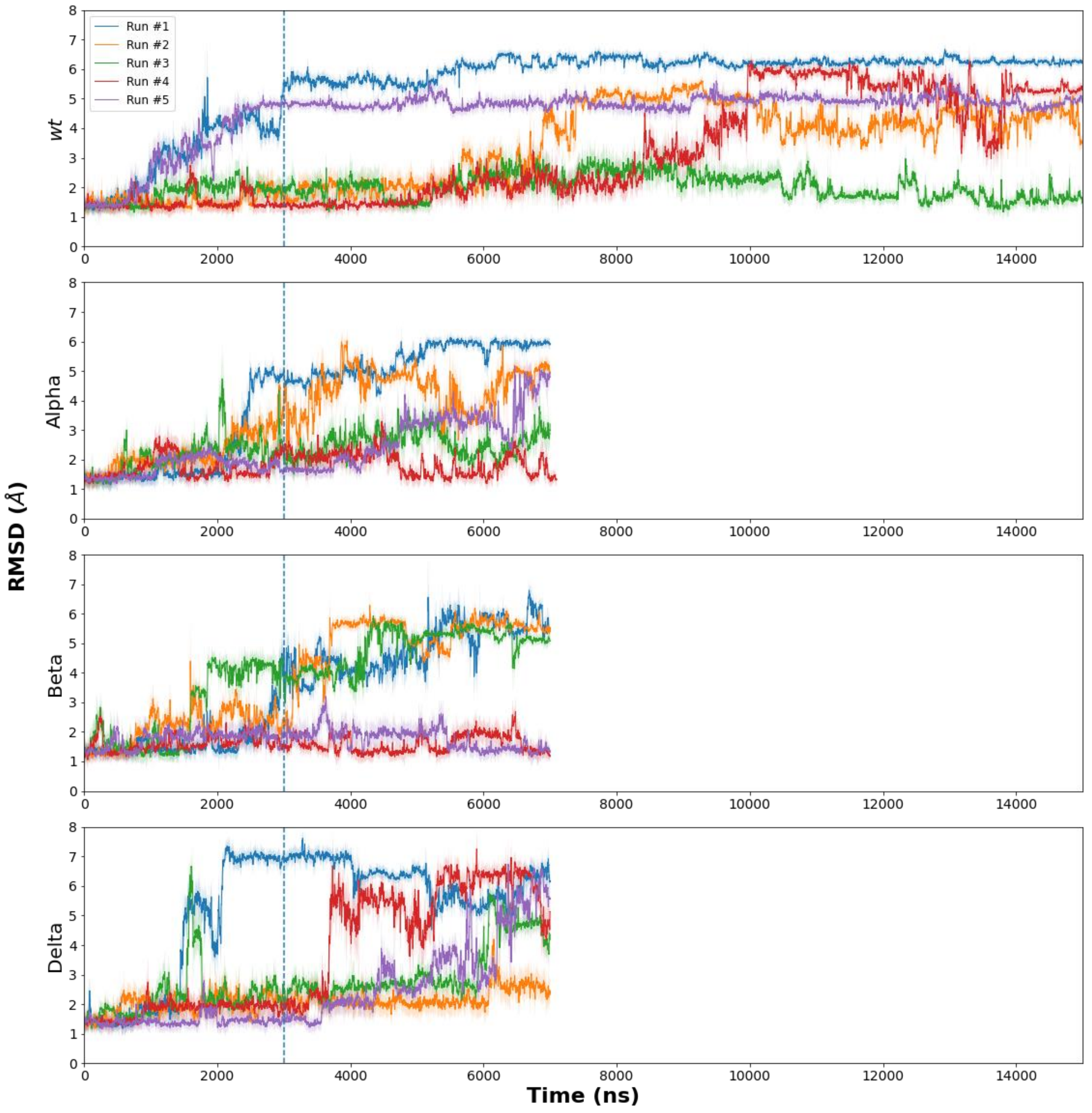


**Figure S12. RBD Cα root-mean-square deviation (RMSD) moving average in solution.** Data shown for the five replicas for each variant tested. Cα were fitted against the RBD X-ray structure from PDB ID: 6M0J. The moving average was calculated using the neighboring 50 frames. The first 3 μs of simulation were used for equilibration (blue dashed line) and the remaining frames were used for further PCA and RIN analysis.

### REFERENCES


(1) Abraham, M. J.; Murtola, T.; Schulz, R.; Páll, S.; Smith, J. C.; Hess, B.; Lindah, E. Gromacs: High Performance Molecular Simulations through Multi-Level Parallelism from Laptops to Supercomputers. *SoftwareX* **2015**, *1–2*, 19–25.

(2) McCallum, M.; Walls, A. C.; Sprouse, K. R.; Bowen, J. E.; Rosen, L.; Dang, H. V.; deMarco, A.; Franko, N.; Tilles, S. W.; Logue, J.; Miranda, M. C.; Ahlrichs, M.; Carter, L.; Snell, G.; Pizzuto, M. S.; Chu, H. Y.; Voorhis, W. C. Van; Corti, D.; Veesler, D. Molecular Basis of Immune Evasion by the Delta and Kappa SARS-CoV-2 Variants. *bioRxiv* **2021**, 2021.08.11.455956.

(3) Tian, F.; Tong, B.; Sun, L.; Shi, S.; Zheng, B.; Wang, Z.; Dong, X.; Zheng, P. N501Y Mutation of Spike Protein in SARS-CoV-2 Strengthens Its Binding to Receptor ACE2. *Elife* **2021**, *10*.

(4) Laffeber, C.; de Koning, K.; Kanaar, R.; Lebbink, J. H. G. Experimental Evidence for Enhanced Receptor Binding by Rapidly Spreading SARS-CoV-2 Variants. *J. Mol. Biol.* **2021**, *433* (15), 167058.

(5) Supasa, P.; Zhou, D.; Dejnirattisai, W.; Liu, C.; Mentzer, A. J.; Ginn, H. M.; Zhao, Y.; Duyvesteyn, H. M. E.; Nutalai, R.; Tuekprakhon, A.; Wang, B.; Paesen, G. C.; Slon-Campos, J.; López-Camacho, C.; Hallis, B.; Coombes, N.; Bewley, K. R.; Charlton, S.; Walter, T. S.; Barnes, E.; Dunachie, S. J.; Skelly, D.; Lumley, S. F.; Baker, N.; Shaik, I.; Humphries, H. E.; Godwin, K.; Gent, N.; Sienkiewicz, A.; Dold, C.; Levin, R.; Dong, T.; Pollard, A. J.; Knight, J. C.; Klenerman, P.; Crook, D.; Lambe, T.; Clutterbuck, E.; Bibi, S.; Flaxman, A.; Bittaye, M.; Belij-Rammerstorfer, S.; Gilbert, S.; Hall, D. R.; Williams, M. A.; Paterson, N. G.; James, W.; Carroll, M. W.; Fry, E. E.; Mongkolsapaya, J.; Ren, J.; Stuart, D. I.; Screaton, G. R. Reduced Neutralization of SARS-CoV-2 B.1.1.7 Variant by Convalescent and Vaccine Sera. *Cell* **2021**, *184* (8), 2201-2211.e7.

(6) Wirnsberger, G.; Monteil, V.; Eaton, B.; Postnikova, E.; Murphy, M.; Braunsfeld, B.; Crozier, I.; Kricek, F.; Niederhöfer, J.; Schwarzböck, A.; Breid, H.; Jimenez, A. S.; Bugajska-Schretter, A.; Dohnal, A.; Ruf, C.; Gugenberger, R.; Hagelkruys, A.; Montserrat, N.; Holbrook, M. R.; Oostenbrink, C.; Shoemaker, R. H.; Mirazimi, A.; Penninger, J. M. Clinical Grade ACE2 as a Universal Agent to Block SARS-CoV-2 Variants. *bioRxiv* **2021**, 2021.09.10.459744.

(7) Souza, A. S. de; Amorim, V. M. de F.; Guardia, G. D. A.; Santos, F. R. C. dos; Santos, F. F. dos; Souza, R. F. de; Juvenal, G. de A.; Huang, Y.; Ge, P.; Jiang, Y.; Paudel, P.; Ulrich, H.; Galante, P. A. F.; Guzzo, C. R. Molecular Dynamics Analysis of Fast-Spreading Severe Acute Respiratory Syndrome Coronavirus 2 Variants and Their Effects in the Interaction with Human Angiotensin-Converting Enzyme 2. *bioRxiv* **2021**, 2021.06.14.448436.

(8) Saville, J. W.; Mannar, D.; Zhu, X.; Srivastava, S. S.; Berezuk, A. M.; Demers, J.-P.; Zhou, S.; Tuttle, K. S.; Sekirov, I.; Kim, A.; Li, W.; Dimitrov, D. S.; Subramaniam, S. Structural and Biochemical Rationale for Enhanced Spike Protein Fitness in Delta and Kappa SARS-CoV-2 Variants. *bioRxiv* **2021**, 2021.09.02.458774.
